# The Crohn’s disease-related AIEC strain LF82 assembles a biofilm-like matrix to protect intracellular microcolonies from phagolysosomal attack

**DOI:** 10.1101/2020.03.31.014175

**Authors:** Victoria Prudent, Gaëlle Demarre, Emilie Vazeille, Maxime Wery, Antinéa Ravet, Nicole Quenech’Du, Julie Dauverd Girault, Marie-Agnès Bringer, Marc Descrimes, Nicolas Barnich, Sylvie Rimsky, Antonin Morillon, Olivier Espéli

## Abstract

Patients with Crohn’s disease exhibit abnormal colonization of the intestine by proteobacteria, and among these bacteria, the adherent invasive *E. coli* (AIEC) family. They are predominant in the mucus, adhere to epithelial cells, colonize them and survive inside macrophages. We recently demonstrated that the acclimation of the AIEC strain LF82 to phagolysosomal stress requires stringent and SOS responses. Such adaptation involves a long lag phase in which many LF82 cells become antibiotic tolerant. Later during infection, they proliferate in vacuoles and form colonies harboring dozens of LF82 bacteria. In the present work, we investigated the mechanism sustaining this phase of growth. We found that intracellular LF82 produced an extrabacterial matrix composed of exopolysaccharides and amyloid fibers that surrounded each individual LF82 cell. This matrix acts as a biofilm and controls the formation of LF82 intracellular bacterial communities (IBCs) inside phagolysosomes for several days post infection. Using genomics assays, we characterized the gene set involved in IBCs formation and revealed the crucial role played by a pathogenicity island presents in the genome of most AIEC strains in this process. Iron capture, by the yersiniabactin system encoded by this pathogenicity island, is essential to form IBC and LF82 survival within macrophages. These results demonstrate that AIEC have developed a sophisticated strategy to establish their replicative niche within macrophages, which might have implications for envisioning future antibacterial strategies for Crohn’s disease.

## Introduction

Patients with Crohn’s disease exhibit abnormal colonization of the intestine by proteobacteria, among which the adherent invasive *E. coli* (AIEC) family has been characterized. AIEC are found in intestinal lesions of patients with inflammatory bowel disease. These bacteria are found mainly in the mucus, they adhere to epithelial cells and colonize them to survive inside macrophages. The mechanisms underlying AIEC adaptation for survival and growth inside macrophages are not yet fully understood. Previous work performed with murine macrophage cell lines has revealed that the prototype AIEC strain LF82 proliferates in a vacuole exhibiting the characteristics of a mature phagolysosome (Bringer *et al*, 2006; Lapaquette *et al*, 2012). In such an environment, AIEC should encounter acidic, oxidative, genotoxic and proteic stress. Screening of genes involved in LF82 fitness within macrophages has revealed that the HtrA, DsbA, or Fis proteins are required for optimum fitness (Bringer *et al*, 2005, 2007; Miquel *et al*, 2010a). These observations confirmed that LF82 encounters stress in phagolysosomes. However, much remains to be learned about the host-pathogen interactions that govern AIEC infection biology. AIEC expresses various virulence factors that might play roles in digestive tract colonization, adherence and invasion of epithelial cell, and intracellular survival in phagolysosomes. The diversity of virulence factors displayed by multiple AIEC strains suggests that members of this pathovar may have evolved different strategies to colonize their hosts (Tawfik *et al*, 2014). However, virulence factors involved in iron capture (the high-pathogenicity island (HPI), *chuABCD, fhuD* or *sitABCD* system) exhibit a high prevalence among AIEC strains (Céspedes *et al*, 2017), suggesting that iron capture is a crucial determinant for AIEC strains to form their ecological niches.

We have recently demonstrated that the stringent response (the bacterial checkpoint involved in dealing with nutrient limitation) and SOS response (the bacterial pathway involved in DNA repair) are critical for AIEC survival and multiplication within macrophages. The triggering of these two stress pathways has direct consequences, creating heterogeneity in the bacterial population with replicating nonreplicating bacteria that contribute respectively to increase population size and tolerate the stress. The nonreplicative population of LF82 tolerates antibiotics when they are phagocytosed by macrophages and for a significant period of time after macrophage death. Such bacteria are called persisters and are suspected to be the cause of relapsing infectious chronic diseases, for instance in tuberculosis or cystic fibrosis.

In addition to persistence, biofilm structures represent another source of antibiotic tolerance. Biofilms are communities of cells attached to surfaces and held together by a self-produced extracellular matrix. The matrix is composed of various extracellular DNA molecules, proteins, and polysaccharides, depending on the bacterial species (Joo & Otto, 2012). Cells in the biofilm state exhibit increased protection against desiccation and harmful substances, including antibiotics and the host immune response molecules (Flemming & Wingender, 2010). Biofilms retain bacteria to diverse surfaces, including, in the case of certain pathogens, tissues and, less frequently, intracellular surfaces. Intracellular biofilms, also called intracellular bacterial communities (IBCs), have been described when uropathogenic *E. coli* (UPEC) invade urothelial cells lining the urinary bladder (Anderson *et al*, 2003). This biofilm allows to escape host defense and UPEC persist despite antibiotic therapy, then replicate rapidly (doubling times of 30-35 minutes) for up to 8 hours to form a loose collection of hundreds of bacteria (Scott *et al*, 2015). At this point, the growth rate slows down, and UPEC develop pods with biofilm-like traits.

In the present work, we investigated the mechanism by which the AIEC pathobionts such as the LF82 strain forms microcolonies inside macrophage phagolysosomes. We found that LF82 microcolonies form IBCs with an extracellular matrix composed of exopolysaccharides and curli fibers surrounding each bacterium. We reveal a list of genes critical in the formation of *E. coli* biofilms that are also required for IBCs and survival in macrophages. Finally, by combining genomic screens (dual RNA-seq and TN-seq) we unveiled an LF82’s pathogenicity island required for IBCs. This pathogenicity island, called HPI, allows iron providing to the IBC. Interestingly our results show, for the first time, the link between iron homeostasis and biofilm matrix production for intracellular pathogen proliferation. HPI is present in many AIEC and pathogen bacteria, suggesting that the strategy developed by LF82 might be common for AIEC and other facultative intracellular pathogens and pathobionts that need to survive in the particular niche of the phagolysosome.

## Results

### The AIEC strain LF82 forms intracellular bacterial communities (IBCs) within phagolysosomes

In the minutes following phagocytosis, the induction of a stringent response blocks bacterial cell division and curbs the expansion of the LF82 population. This 6 to 10 h step can lead to the formation of LF82 persisters (Demarre *et al*, 2019). When LF82 resumes growth, the expansion of the population is dependent on the ability of the bacteria to repair lesions (e.g. DNA lesions). Bacteria involved in this multiplication stage divide 4 - 6 times in 10 h. This growth phase leads frequently to the formation of vacuoles containing more than 20 bacteria within human THP1 macrophages derived from monocytes and up to hundreds of bacteria in murine Raw macrophage cell lines (Figure 1A and 1B). To discriminate whether the formation of large vacuoles corresponded to only the clonal multiplication of one or few phagocytic bacteria or, alternatively, to the fusion of different vacuoles, Raw macrophages were infected successively (1 h interval) with GFP-tagged LF82 and mCherry-tagged LF82 (Figure 1C). One hour post infection (P.I.) with LF82-mCherry, most vacuoles contained only one type of LF82 (either green or red), while twenty-four hours P.I., a small proportion of vacuoles presented both types of LF82 (Figure 1C, low panel). Surprisingly, we observed that red and green LF82 formed clonal sectors within vacuoles. These observations suggest that the formation of large vacuoles is not the consequence of multiple fusion events and that LF82 is not free to move within a vacuole. To directly measure the ability of LF82 to move inside a vacuole, we used fluorescence recovery after photobleaching (FRAP) experiments. Bleaching of several spots inside large vacuoles did not lead to recovery of the fluorescence and demonstrated strong adherence to LF82 (Figure 1D and 1E). We deduced that limited movement of LF82 promoted the formation of colonies within phagolysosomes, akin to the IBCs observed for UPEC within bladder cells (Anderson *et al*, 2003).

**Figure 1:**
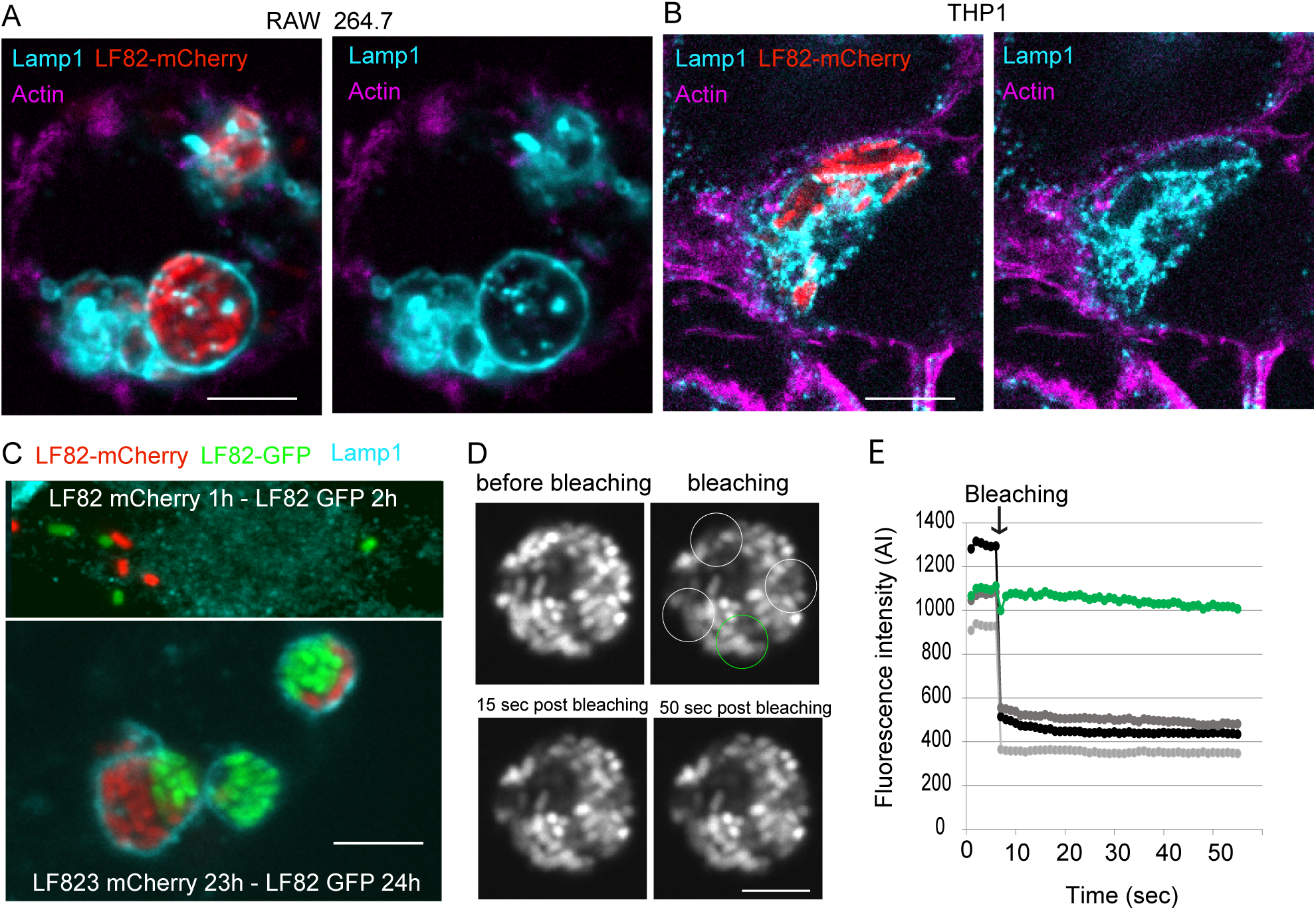
Intracellular LF82 forms intracellular bacterial communities inside phagolysosomes. A) Imaging of the phagolysosomes (antibody for Lamp1, cyan) of Raw macrophages (actin labeling with phalloidin, magenta) infected by LF82-mCherry (red) at an MOI of 30 at 24 h P.I. B) Imaging of the phagolysosomes (antibody for Lamp1, cyan) of THP1 macrophages (actin labeling with phalloidin, magenta) infected by LF82-mCherry (red) at an MOI of 30 at 24 h P.I. The scale bar is 5 µm. C) Coinfection experiment revealing LF82 clonal clusters formed by the fusion of different vacuoles within phagolysosomes. Raw macrophages were infected first with LF82-mCherry, treated with gentamycin for 1 h to remove free LF82-mCherry and subsequently infected with LF82-GFP. At 24 h P.I., phagolysosomes were labeled with the Lamp1 antibody (cyan) and imaged. The scale bar is 5 µm. D) FRAP experiments on LF82-GFP phagolysosomes at 24 h P.I. White circles outline the bleached regions, and green circles outline the control region. The scale bar is 5 µm. E) Fluorescence quantification of the FRAP experiment presented in D.

### Transcriptomic analysis of LF82 survival within macrophages

We performed a dual RNA-seq experiment of THP1 macrophages infected by LF82 at 1 and 6h P.I. to characterize the transcriptional program corresponding to the formation of IBCs. Experiments were performed in duplicate; about 4 million bacterial and 300 million human reads per sample were collected (Table S1) and DEseq P-values were calculated for each gene (Figure 2A and 2B). We first analyzed the bacterial transcriptome. The transcriptomic response of LF82 during macrophage infection was considerable; the expression of 700 and 1000 genes was significantly changed at 1 and 6 h P.I., respectively (Figure 2A and Table S2). At 1h P.I., a majority of the genes were downregulated, while at the 6h, both upregulated and downregulated genes were detected. Genes involved in the responses to the toxic phagolysosome’s environment were among the most significantly upregulated while genes involved in the carbon metabolism and bacterial motility were the most significantly down regulated (Figure 2A). COG analysis confirmed this observation, revealing that the main upregulated pathways were the response to external stimuli (particularly pH variation and SOS response), biofilm regulation, cell communication and organophosphate metabolic processes (Table S3). The main downregulated pathways were metabolic pathways, including energetic metabolism, nucleoside phosphate metabolism, and carbohydrate metabolism (Table S3). To get a global view of the transcriptomic response, we analyzed the most significantly changed 500 genes at 1 h (Supplementary Figure S1A) and 6 h P.I. (Figure 2C). We grouped genes in 22 categories based on available annotations. Some categories were densely populated (Central metabolism, for example, present nearly 100 genes with significant mRNA fold change at 1 h or 6 h P.I.) and some contained only few representatives (3 efflux pumps). This analysis gave an image of the growth conditions encountered by AIEC LF82 within macrophages. It confirmed the presence of acid, oxidative and genotoxic stresses, and showed: i) the alteration of the bacterial envelope (membrane, cell wall and periplasm), ii) the reduction of protein biosynthesis and cell cycle activity, presumably linked to stringent response induction (Demarre *et al*, 2019), iii) the switch from motile to biofilm behavior and, iv) the global reduction of catabolic and energetic metabolism. We used data available for *E. coli* K12 in the Regulon Database (Gama-Castro *et al*, 2016) to identify the key responses. The Regulon Database provides insights into the relationships of 7031 pairs of genes and regulators (regulators involved in these transcriptomic transcription factors (TFs), sigma factors, sRNAs); among them, the expression of one or two partners was changed in 1400 couples at 6h P.I. The box plot presented in Supplementary Figure S1B illustrates these changes for upregulated genes (green) and downregulated genes (red) for each regulon. Some regulons were mostly (eventually entirely) upregulated or downregulated, but many of them presented both upregulated and downregulated genes (Supplementary Figure S1B). Among the upregulated regulons, those associated with the response to acidic pH are largely overrepresented. These regulons are controlled by the EvgA, GadE, GadX, GadW, YdeO, PhoP, and BluR regulators and involve 130 genes. Consistent with our previous observations, genes involved in the response to acidic pH are highly overexpressed as early as 1h P.I. (Supplementary Figure S1C). Members of the second family of of upregulated regulons govern the bacterial growth mode (cell division, LPS, capsule or biofilm determinants). The BluR, CsgD, Dan, DicA, FliZ, McaS, MlrA, MqsA, NhaR, PhoB, OmpR and RcsAB regulators, control genes in these pathways. Among the genes controlled by these TFs, we observed, upon entry into the macrophage, a general upregulation of the genes annotated for biofilms and adhesion (Supplementary Figure S1C). Consistent with this change in the LF82 growth, upon entry into the macrophage, we observed a severe downregulation of the FlhDC regulon, which contains motility genes (Supplementary Figure S1C). Biofilm formation frequently corresponds to a switch toward a slower metabolism, which is exactly what the RNA-seq data suggested, with downregulation of genes involved in glycolysis, ATP production and purine/pyrimidine metabolism (Supplementary Figure S1C). We mapped RNA-seq results on the KEGG ko2026 pathway for biofilm formation of commensal E. coli (Figure 2D). This map suggests that the lack of nutrient, perhaps some amino acids, induces stringent response and consequently represses the expression of FlhDC, therefore flagellar assembly is stopped and bacteria become non-motile. At the same time acid, osmolarity and envelope stresses lead to the induction of *csgD* via OmpR and RpoS. CsgD induces the expression of the operons encoding curli fibers. Quorum sensing, via the BarA/UvrY two component system, the LuxS/LsrR system and the CsrA repression will lead to the expression of the operons involved in the production of poly-N-acetyl glucosamine and colanic acid. On this map, the role played by ci-di-GMP is not easily understandable; diguanylate cyclase genes involved in the production of ci-di-GMP, are either downregulated (*dgcQ* and *dgcM*) or upregulated (*adrA*) and cyclic di-GMP phosphodiesterases *pdeR* (*gmr*) is upregulated while *pdeH* is downregulated. Genes involved in the production of cellulose are repressed; the available literature does not allow to explain this observation. These transcriptomic observations strongly suggest that LF82 switches from a planktonic to a biofilm mode of growth once within macrophage and suggest that an extracellular matrix is responsible for the observed adherence and clonal growth within IBCs.

**Figure 2:**
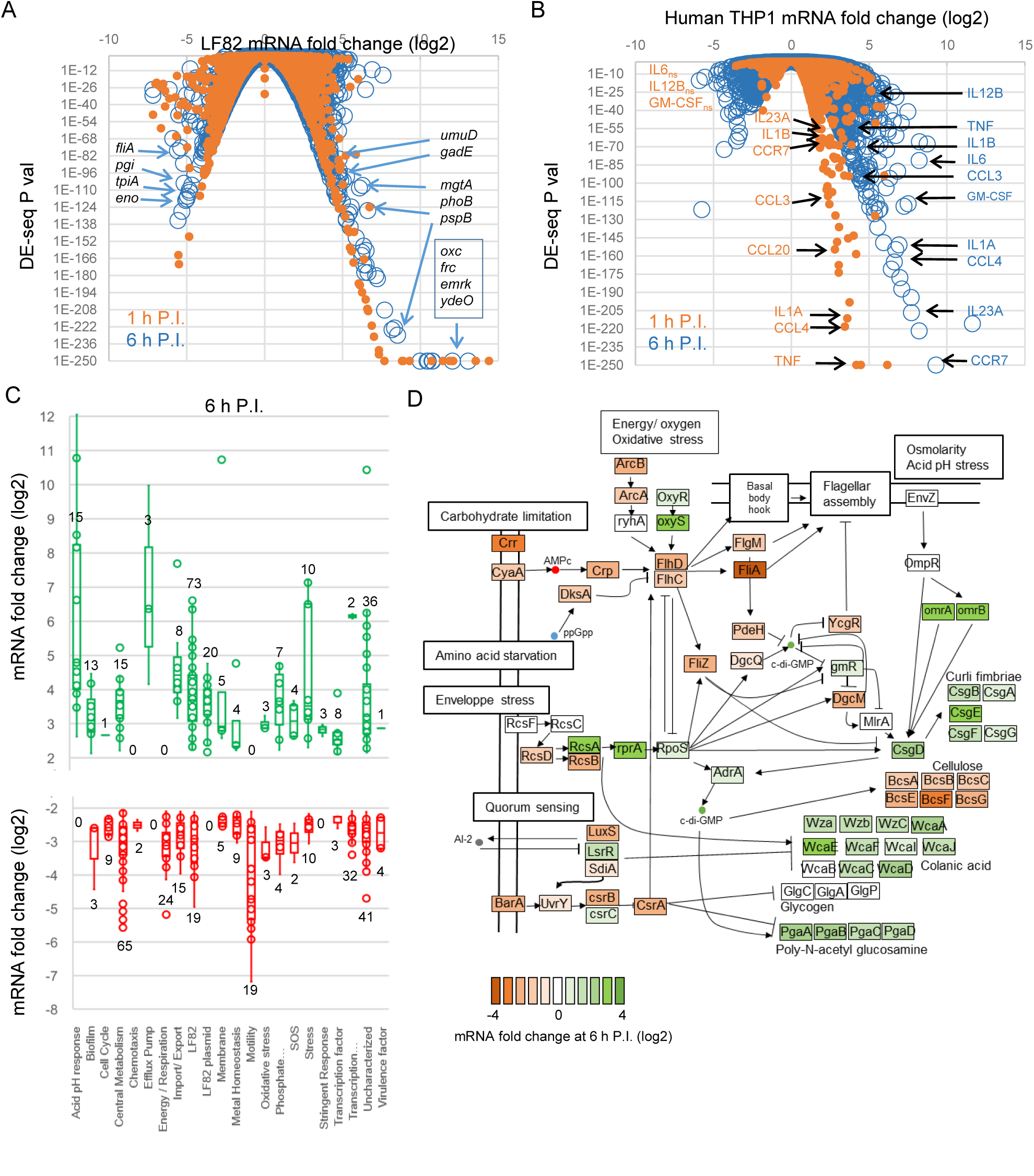
Transcriptomic analysis of macrophage infection by LF82. A) Expression data for the LF82 genes from RNA-seq experiments at 1 h and 6 h P.I. in THP1 macrophages. Selected genes with highly significant expression changes at 6 h P.I. are noted. B) Expression data for the human genes from RNA-seq experiments at 1 h and 6 h P.I. in THP1 macrophages. Gene markers of the M1 pro inflammatory macrophage phenotype are noted. When mRNA fold changes are not significant they are noted with an “ns” index. Orange labeling (1 h P.I.) and blue labeling (6 h P.I.). C) Analysis of the 500 LF82 genes presenting the most significantly changed mRNA fold at 6h P.I.. The data for the 1 h P.I. time point are presented on Supplementary Figure S1. Gene annotations were manually curated to define 22 categories: Acid pH response, Biofilm matrix and regulation, Cell cycle, Central metabolism (carbohydrates, nucleotides, amino acids), Chemotaxis, Efflux pumps, Energy and respiratory metabolism, Import and export of nutrients, LF82 genes without homologue in the E. coli K12 genome, LF82 genes encoded by the mega plasmid pLF82, Membrane and envelope components, Metal homeostasis, oxidative stress, Phosphate homeostasis, SOS response, Diverse stress response (including cold shock proteins and phage shock proteins), Stringent response, Transcription factors, uncharacterized genes and putative virulence factors (presenting an homolog in the K12 genome). Red (upregulated genes) and green (downregulated genes). Numbers on the graph indicate the number of genes in the given category. D) Annotation of the biofilm formation pathways adapted from KEGG ko2026 according to RNA-seq data at 6h P.I. vs planktonic exponential growth.

#### Transcriptomic response of the macrophages to AIEC infection

We analyzed the Dual RNA-seq data to monitor macrophage response to LF82 infection. LF82 infection significantly impacted the THP1 macrophage transcriptome (Figure 2B, Table S6). As expected, we observed the induction of the antimicrobial humoral immune response (the CXCL1, CXCL2, CXLC3, IL1 and IL8 genes were among the most significantly affected genes); the inflammatory response (TNF, CCR7, NFKBIA, OSM, CCL3L3, NFBIZ, PTSG2); TNF and the response to its production (the CCL1, CCL3, CCL3 and CCL20 genes); and the lymphocyte activation pathway (CD80, IL23A). The response is stronger (i.e. more genes significantly changed, increased number of COG represented in the list of significantly changed genes and higher mRNA fold ratio) at 6h P.I. compared to 1 h. P.I. Upon infection macrophages become polarized, this is frequently illustrated by the presence of two phenotypes: M1 like macrophages that are pro inflammatory and M2 like macrophage that are anti inflammatory. Recent work demonstrated that salmonella is able to drive polarization of mice BMM macrophages toward an M2 more permissive state (Stapels *et al*, 2018). In human, the separating line between M1 and M2 like macrophages is rather represented by a continuum where boundaries are still unclear. However several transcriptomic markers were attributed to M1 (IL12B, CCR7, IL1A, IL1B, IL23A, TNF, CCL3, CCL4, GM-CSF) and M2 (GM-CSF, IL4, IL10, IL13, CCL1, CCl2, CCL17, CCL18, CCL22, CCL23, CCL24, CCL26, IL1R, TGFB) (Atri *et al*, 2018). RNA-seq data showed that most M1 markers are significantly overexpressed upon LF82 infection (Figure 2B) while none of the M2 markers were upregulated (Table S6). This confirmed that LF82 encounter stressful environment during its first hour of survival within pro inflammatory macrophages.

### IBCs are structured by an extracellular matrix

Using fluorescent lectins to visualize the exopolysaccharide matrix, we detected Wheat germ agglutinin (WGA) and soybean agglutinin (SBA) around bacteria in the phagolysosomes (Figure 3A and Supplementary Figure S2A). Labeling of WGA and SBA revealed that both form envelopes surrounding each bacteria. WGA labeling was weak at 1h P.I. while it was clearly detected in Raw 264.6, THP1 and HDMM at 24h P.I. (Supplementary Figure S2B-D). In addition, we also frequently observed a strong labeling of bacterial periphery that colocalizated with Lamp1. WGA and SBA were also detected in many other organelles of the macrophage including Golgi apparatus that contains glycosylated proteins and lipids. To challenge the specificity of the WGA staining, we performed coinfection with a K12 C600 *E. coli* strain that is not as efficient as LF82 to proliferate within macrophages, making rare occasional foci consisting of several bacteria (Figure 3B). In this assay, WGA labeling was strongly reduced in the C600 vacuoles compared to the LF82 vacuoles (Figure 3B and 3C). We used STED superresolution microscopy to refine our analysis of the WGA staining in WT LF82 and LF82 lacking the regulator *csgD* and the *pgaABCD* operon encoding the exopolysaccharide component of the matrix. WGA labeling around bacteria inside the phagolysosomes was observed for the WT strain while no WGA labeling was detected around LF82 Δ*csgD* and LF82Δ*pgaABCD* (Figure 3D). All together, these observations confirmed that exopolysaccharides observed around LF82 are from bacterial origin. The absence of an exopolysaccharide matrix in LF82Δ*pgaABCD* was expected because this operon directly encodes the enzymes dedicated to the synthesis and export of exopolysaccharide; however, the connections between CsgD and the exopolysaccharide matrix have not been documented in commensal *E. coli* (Parker *et al*, 2017). Prolonged infection experiments showed that WGA labeling was present in the IBC formed by LF82 up to 72h P.I., but absent in LF82 Δ*csgD.* This suggest that exopolysaccharide production is abolished rather than just delayed in the *csgD* mutant (Supplementary Figure S2E). Thus, we also concluded that CsgD is an important factor to build LF82-specific matrix within macrophage. Because CsgD is the main controller of the production of curli fibers, we analyzed curli by immunostaining with antibodies against CsgA or CsgB (Zhou *et al*, 2013); unfortunately, we could not observe labeling inside macrophages with antibodies (Figure 3E). Therefore, we tested the antibodies on bacteria recovered from lysed macrophages 24h P.I. and LF82 showed strong CsgA labeling at its periphery, while C600 presented much weaker labeling (Figure 3F and 3G) and the LF82 *csgD* mutant did not show any labeling (Figure 3H). The genes allowing the biosynthesis and export of colanic acid, another exopolysaccharide, were also expressed in the phagolysosomes. However, we did not detect any specific labeling of LF82 vacuoles with Concanavalin A (ConA) or Peanut Agglutinin (PNA) that reveals β galactosides present in colanic acid (Supplementary Figure S3). The genes encoding the protein responsible for cellulose biosynthesis and export were downregulated in the macrophage (Figure 2D) suggesting that cellulose is not involved in LF82 IBC. Our results demonstrate that LF82 produce a biofilm-like matrix containing at least poly-β-1,6-*N*-acetyl-D-glucosamine exopolysaccharide and amyloid fibrils composed of curlin to organize IBCs.

**Figure 3:**
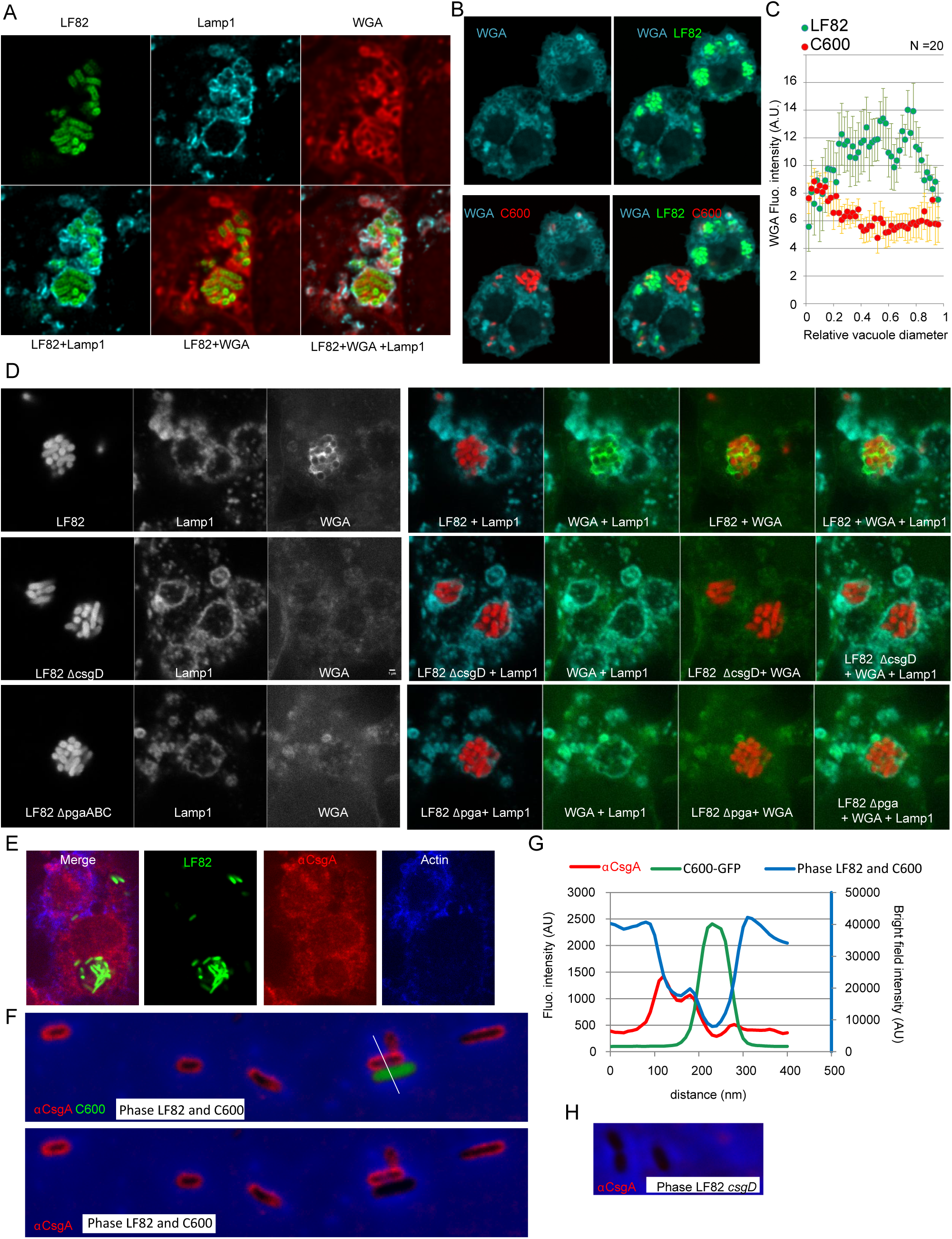
Intracellular LF82 IBCs present a biofilm-like matrix. A) Imaging of extracellular polysaccharide structures revealed by wheat germ agglutinin (WGA) labeling around LF82-GFP (green) inside phagolysosomes (antibody for Lamp1, cyan) of Raw macrophages. MOI of 30, 24 h P.I. B) At 24 h P. I. polysaccharide structures are observed inside phagolysosomes containing LF82-GFP but not inside phagolysosomes containing only C600-mCherry bacteria. C) Quantification of the WGA fluorescence intensity along the vacuole diameter of raw macrophages infected by LF82 or C600 for 24 h P.I.. Values represent median +/- SD of 20 vacuoles. D) STED superresolution imaging of the extracellular polysaccharide structures revealed by WGA staining around LF82 IBCs formed by the WT and the *csgD* and *pgaABC* mutants. E) Imaging of curli fibers by conventional immune labeling with the antibody for CsgA (red) of fixed and permeabilized macrophages infected by LF82 F) Imaging of curli fibers (antibody for CsgA, red) formed around LF82 during macrophage infection. Macrophages infected with LF82 and C600-GFP were lysed with Triton 1x at 24 h P.I. The lysate was immediately fixed with paraformaldehyde, spread on agarose pads and imaged. G) Quantification of the fluorescence intensities along the white line drawn in D. H) Imaging of curli fibers (antibody for CsgA, red) in the LF82 *csgD* strain, the experiment was performed as described in F fir the WT strain.

### The extracellular matrix affects IBC’s size and LF82 survival within macrophages

To characterize better the mechanism by which biofilm-like matrix involved in IBC, we constructed deletion mutants lacking genes controlling extracellular matrix production (*csgD, adrA, dgcE, pdeH, ompR, rcsCBD*), genes encoding for structural components of the matrix (*pgaA, wza, waaWVL, bcsB, glgCAP*), genes involved in surface adhesion (*fimH)* and quorum sensing (*qseBC*). We tested the ability of these mutants to form biofilms on the abiotic surface of 96-well polystyrene plates. Most single mutants presented a clear defect to form biofilms (Supplementary Figure S4); although they maintained a higher biofilm formation than the C600 laboratory strain. We noted that the *pdeH* and *fimH* mutants were severely affected, confirming that cyclic di-GMP regulation and adhesion are required for the formation of the biofilm. We then tested some of these mutants for their ability to survive for 24h within Raw 264.7 macrophages, THP1 macrophages and human monocyte-derived macrophages (HMDMs) from blood (Figure 4A-4C). In Raw macrophages, viability was reduced by 20 to 60 % for most of the mutants. In THP1 macrophages, viability was reduced but to a lesser extent. Finally, in the HMDMs, the viability of the *csgD* mutant was also significantly reduced. One important aspect of LF82 survival within macrophages is the formation of nongrowing bacteria immediately upon infection and control of the stringent response (Demarre *et al.*, 2017). IBC are formed during the later replicative phase, starting 10h P.I., we observed that only 4 −10% of the vacuoles contained more than 16 bacteria at 24h P.I.. In addition, many LF82 were included in small vacuoles with less than 8 bacteria (Figure 4 D and E). In contrast, using the *rcsBD, pgaA, csgD, waaWVL, wza* mutants we observed that, in average, vacuoles contain less bacteria. This suggests that when the biofilm matrix is altered the proportion of LF82 able to form or to maintain large IBC structures is affected. Altogether, these results show that IBCs constitute an important determinant of the LF82 infection program.

**Figure 4:**
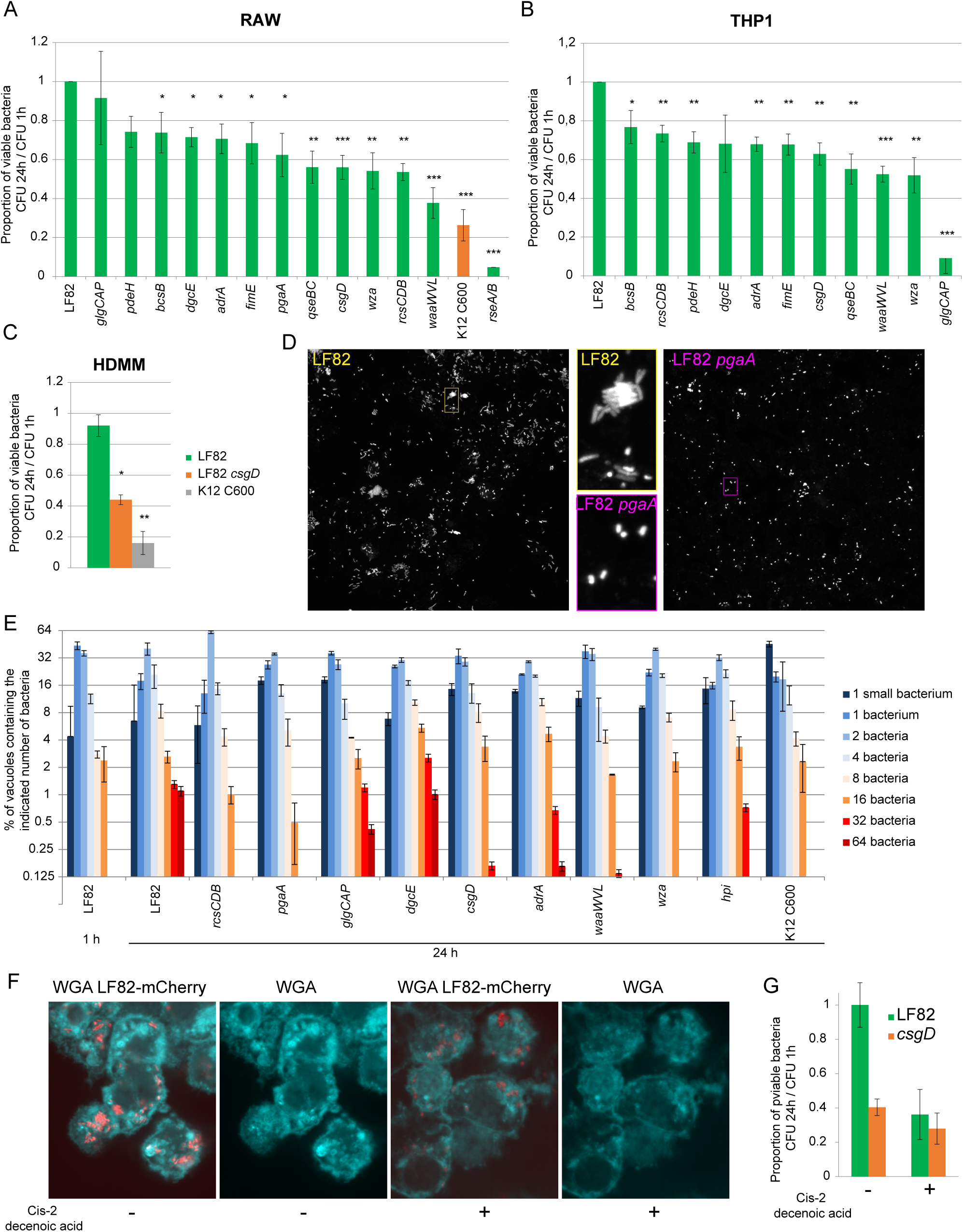
Deletions of the genes involved in the synthesis of the extracellular matrix curb LF82 survival and antibiotic tolerance. A) Proportion of viable bacteria at 24 h P.I. in Raw macrophages in comparison to those present at 1 h. LF82 and LF82 deletion mutants (green) and K12-C600 (orange) were infected at an MOI of 30. Values represent the average of 3 to 7 experiments ± SD. The data were analyzed using Student’s *t* test: * *P* <0.05, ** *P* <0.01, *** *P* <0.001, **** *P* <0.0001. B) Proportion of viable bacteria at 24 h P.I. of THP1 macrophages in comparison to those present at 1 h. The data were analyzed as in A. C) Proportion of viable LF82, LF82 csgD and K12-C600 bacteria at 24 h P.I. in human derived macrophages from blood monocytes (HDMMs) in comparison to those present at 1 h. The data were analyzed as in A. D) Imaging of the IBCs formed at 24 h P.I. with LF82-mCherry and the LF82 *pgaA* mutant. E) Distribution of IBC’s size in the population of the WT LF82 strain and the *rcsBD, pgaA, glgCAP, dgcE, csgD, adrA, waaWVL, wza, hpi* mutants and the K12 C600 strain. F) Inhibition of exopolysaccharide matrix synthesis by the addition of cis-2-decenoic acid at 6 h P.I.; imaging was performed at 24 h P.I. G) CFU of LF82 and the LF82 *csgD* mutant in the presence of cis-2-decenoic acid in the medium.

### Chemical inhibition of biofilm matrix formation affects the formation of IBC and LF82 survival

To further test the impact of IBCs on the survival of LF82, we used cis-2-decenoic acid, a well-described inhibitor of biofilm formation and stability. We observed a reduction in WGA intensity around LF82, suggesting a reduction in the amount of exopolysaccharide in the matrix (Figure 4F), and concomitantly, we observed a reduction in LF82 survival within macrophages (Figure 4G). The survival of the LF82 *csgD* mutant was not affected by cis-2-decenoic acid, confirming that the drug targets the biofilm matrix. Altogether, our results demonstrate that the IBC structure formed within phagolysosomes is beneficial for LF82 survival in macrophages.

### LF82 genes involved in macrophage survival

To monitor the formation of the extracellular matrix and design mutants for viability assays, we used the extensive knowledge available in the literature on biofilms of laboratory *E. coli* strains. However, we observed a clear difference in matrix formation and survival between LF82 and K12. These observations suggest that some LF82 specific genes or mutations might be responsible for its higher biofilm formation capacity and its ability to form IBC within macrophages. The genome of LF82 contains 925 genes that are not present in the genome of *E. coli* MG1655, among which 122 are encoded by the pLF82 plasmid. Many LF82 genes were upregulated during infection, 172 (26 encoded by pLF82 genes) of these genes were upregulated 1 h P.I., and 205 (46 encoded by pLF82) were upregulated 6 h P.I.. Only 25 (3 from pLF82) and 42 (2 from pLF82) LF82 genes were downregulated 1 h and 6 h P.I., respectively (Figure 2C and Table S1). This suggests that some of the LF82 genes might be selected for intracellular growth and eventually for adaptation to macrophages. To shorten this list of putative candidates, we performed transposon mutagenesis and analysis by whole genome sequencing, Tn-seq (Figure 5A). Knowing that, in average, vacuoles contain 4 - 5 times more LF82 in the Raw 267.4 macrophages compared to the THP1 cell line, we performed this experiment with the Raw cell line. Preliminary experiments suggested that one round of 24 h of infection is not enough to obtain a clear picture of the important genes for the infection. We, therefore, performed three rounds of successive 24 h infections; we allowed a 24 h in LB regrowth step in between each infection. The density of transposon insertion per genes was measured for the population that went through macrophages (3 × 24 h in macrophage + 3 × 24 h in LB) and kept in LB (3× 24 h in LB) (Figure 5A). Our two replicates of the whole experiment were in good agreement (Figure 5A). We used fold change of the average insertion density for further analysis (Figure 5B and 5C). Approximately 180 genes were identified in that present a normalized insertion density > 2 in the library (i.e. genes that are not essential for LF82 in LB) and a reduced fold change in the selection experiment (Figure 5B, Table S4). This group is significantly enriched in transcription factors and phospholipid transport (Table S5). The veracity of our analysis pipeline was supported by findings at the top of the hit list of presence of previously validated important genes important for LF82 survival within macrophages (*dksA, rpoS, slyB, degP, pspB-C-F, phoP*, Figure 5B and Table S4), and important pathways such as DNA repair (*uvrABC, recO* and *ugd*) or envelope homeostasis (*nlpB-C-D-F, pspB-C-F, degP*). It suggested that many Tn-seq hits were genuinely important genes for LF82 survival and growth within macrophages. We observed only weak Tn-seq signals for the genes encoding for proteins involved in the formation of the biofilm matrix (Table S4). However, we observed strong essentiality signals for regulators of the biofilm pathway (*rpoS, dksA, barA, uvrY, rcsB, proQ, comR, bhsA*) (Figure 5B). The lack of genes involved in the building of the exopolysaccharide and curli matrix suggest a certain redundancy in between matrix components or non-cellular autonomous effects allowing mutated bacteria to be protected by the matrix produced by WT LF82 present in the same vacuole. By contrast, the strong selective pressure imposed on the regulators of the biofilm pathway suggests that optimum growth within the IBC is not solely linked to the production of the matrix but also to the expression of a dedicated transcription program. We combined RNA-seq and Tn-seq data to screen for LF82 specific genes putatively involved in IBC growth or survival (Figure 5C). Three gene clusters emerged from this analysis: the pathogenicity island II (also called HPI, in other pathogens), a putative type 6 secretion system (T6SS) and a putative carbohydrate’s metabolism gene cluster. HPI expression was induced at 6h P.I. but not at 1h P.I. (Figure 5D), which correlated with the appearance of robust exopolysaccharide staining (Supplementary Figure S2). In contrast, overexpression of the phosphoglucoside and T6SS clusters was observed as early as 1h P.I. (Supplementary Figure S5). We, therefore, focused our analysis on the HPI, the function of which in *Yersinia pestis* has been well characterized. HPI allows the production of a siderophore called Yersiniabactin, its export and its reentry into the bacteria when iron has been captured. Yersiniabactin is a small peptide synthesized by non-ribosomal peptide synthases encoded on the island (*irp1* and *irp2* genes). Our Tn-seq data revealed that insertions in *ybtX, irp1, irp2, ybtE* and *fyuA* genes were counter selected in the first round of infection but not when LF82 was subjected to three rounds of selection (Figure 5D). In contrast, the *ybtS, ybtP, ybtQ, ybtT* and *ybtU* genes were still under selective pressure after three rounds of infections (Figure 5D). A possible explanation for these observations is that Yersiniabactin production and import is advantageous during infection but might be detrimental during the recovery periods in LB between infections.

**Figure 5:**
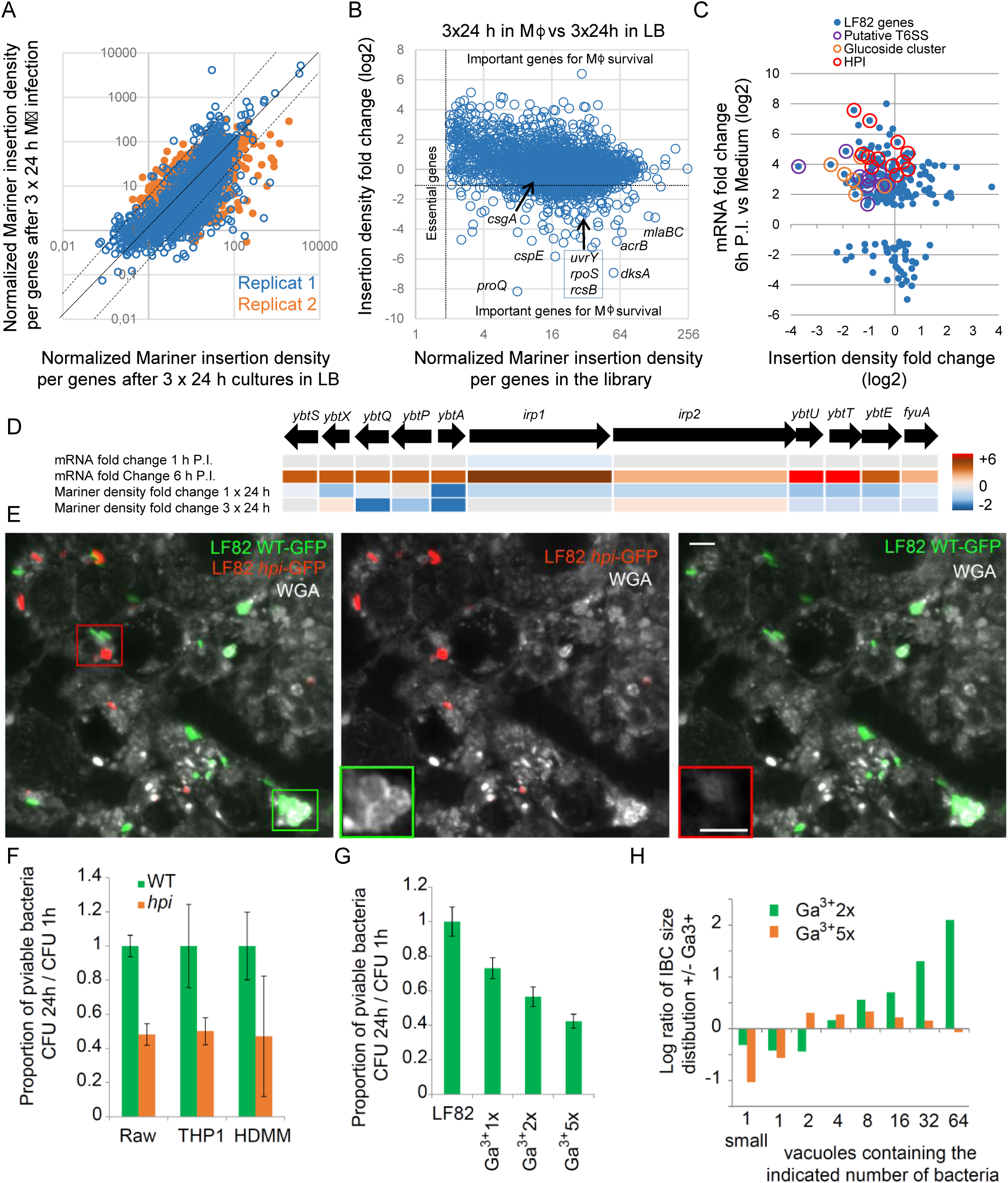
The high-pathogenicity island (HPI) contributes to IBC formation. **A)** Scatter plot representing Tn-seq replicates. B) Scatter plot representing Tn-seq data. Genes presenting a high insertion density in the library (X axis) are under low selective pressure in LB and genes presenting low insertion density fold change (Y axis) are important for survival in the macrophage context. Selected important genes are annotated. C) Scatter plot representing RNA-seq and Tn-seq data for LF82-specific genes (not present in the genome of *E. coli* K12). Only genes with a DE-seq P-value <10^−10^ are represented. Genes highlighted with a orange circle are members of the putative sugar metabolism cluster localized at 1566633-1572808 bp, red circles highlight genes from the pathogenicity island PAI-II or high-pathogenicity island (HPI; 2009366-2039595 bp), and purple circles highlight genes from a putative T6SS (1404073-1517973 bp). D) Summary of RNA-seq and Tn-seq data for the HPI genes. E) Imaging of LF82-GFP and LF82Δ*hpi* – mCherry co-infection 24h P.I.. Exopolysaccharide matrix was stained with lectin WGA. Images are Z projection of the maximum intensity of 10 planes spaced by 400nm. Insets correspond to the WGA labeling at the best focal plane for WT (green) and *hpi* mutant (red). G) Proportion of viable bacteria at 24 h P.I. in Raw, THP1 and HDMM macrophages in comparison to those present at 1 h. LF82 (green) and LF82 *hpi* deletion mutant (orange).F) Proportion of viable bacteria at 24 h P.I. in Raw, macrophages in comparison to those present at 1 h in the presence of Ga^3+^ in the medium. H) Distribution of IBC’s size in the population of the WT LF82 strain in the presence of Ga^3+^ (N=500).

### Iron scavenging by yersinibactin determines LF82 fate within macrophage

HPI is composed of two divergent open reading frames controlled by a single AraC-like regulator (LF82_p299). The *ybtS, ybtX, ybtQ*, and *ybtP* genes are on the Crick strand, and the encoding *ybtA* (encoding the regulator), *irp1, irp2, ybtU, ybtT, ybtE* and *fyuA* (encoding the the outer membrane receptor of Fe^3+^-yersiniabactin) are on the Watson strand. We obtained a mutant in which the first part of the island from *ybtS* to *irp1* was deleted (called *hpi* mutant). This deletion removes the three main promoters of the HPI located around *ybtA*. We observed that the number and size of replicative foci were reduced with the *hpi* mutant strain (Figure 4E and 5E), and the remaining foci did not present WGA labeling for exopolysaccharides around bacteria even if they have been phagocyted by the same macrophage as a WT LF82 (Figure 5E). Viability in Raw, THP1 and HMDMs is affected by the deletion present in *hpi* mutant (Figure 5F). Gallium salts are competitors of iron for siderophores, because of this property they have been successfully used as antibiotics against pathogenic *E. coli* (Choi *et al*, 2019). As expected, the addition of Ga^3+^ in the medium reduced the proportion of viable LF82 at 24h P.I. (Figure 5G). Interestingly, we observed that the proportion of large IBCs increased in the presence of moderate dose of Ga^3+^ suggesting that bacteria from IBC get a privileged access to iron or a protection from the toxic effects of Ga^3+^ (Figure 5H). Therefore, we postulate that yersiniabactin production might be higher inside IBCs compared to single cells.

### Regulation of Iron capture by LF82 within macrophage

To monitor HPI expression at the single cell level, we constructed a plasmid reporting the expression of the HPI promoter with an unstable GFP (P-HPI-GFP*). The expression of HPI started at 6h P.I., as expected from the RNA-seq data, and was evident at 24h P.I. (Figure 6A). At 24h P.I., HPI expression was bimodal, with bacterial foci strongly expressing HPI while the other foci did not express HPI (Figure 6A-6B). Live imaging confirmed this bimodality with an expression starting in some IBC at 4h and reaching a plateau at 10h P.I. while some other bacterial foci remained repressed for the entire kinetics (Supplementary Figure S6). HPI expression was less frequent in the LF82 *csgD* (Figure 6A and 6C), *adrA, bcsB, qseB* and *glgCAP* mutants (Figure 6D), suggesting that the presence of the biofilm matrix is important for HPI expression. In *Yersinia pestis*, HPI expression is controlled by YbtA, a transcriptional activator, and the ferric uptake regulator Fur (Anisimov *et al*, 2005; Gao *et al*, 2008). Fur also controls the expression of other iron capture systems such as the enterobactin, present in the core genome of enterobacteria. LF82 genome present 5 iron capture systems: the enterobactin system, the *sitABCD* system, the *Chu* haem capture system, a putative iron transpoter system and the HPI. Our RNA-seq experiments showed that inside macrophages, most iron capture systems were repressed, except the one encoded by HPI, which was highly overexpressed (Figure 6E). This observation suggests that the a particular regulation of the HPI is at play within IBCs and that the Yersiniabactin system, in particular, is adequate for iron capture in the specific context of the IBCs of the AIEC LF82 strain within phagolysosomes.

**Figure 6:**
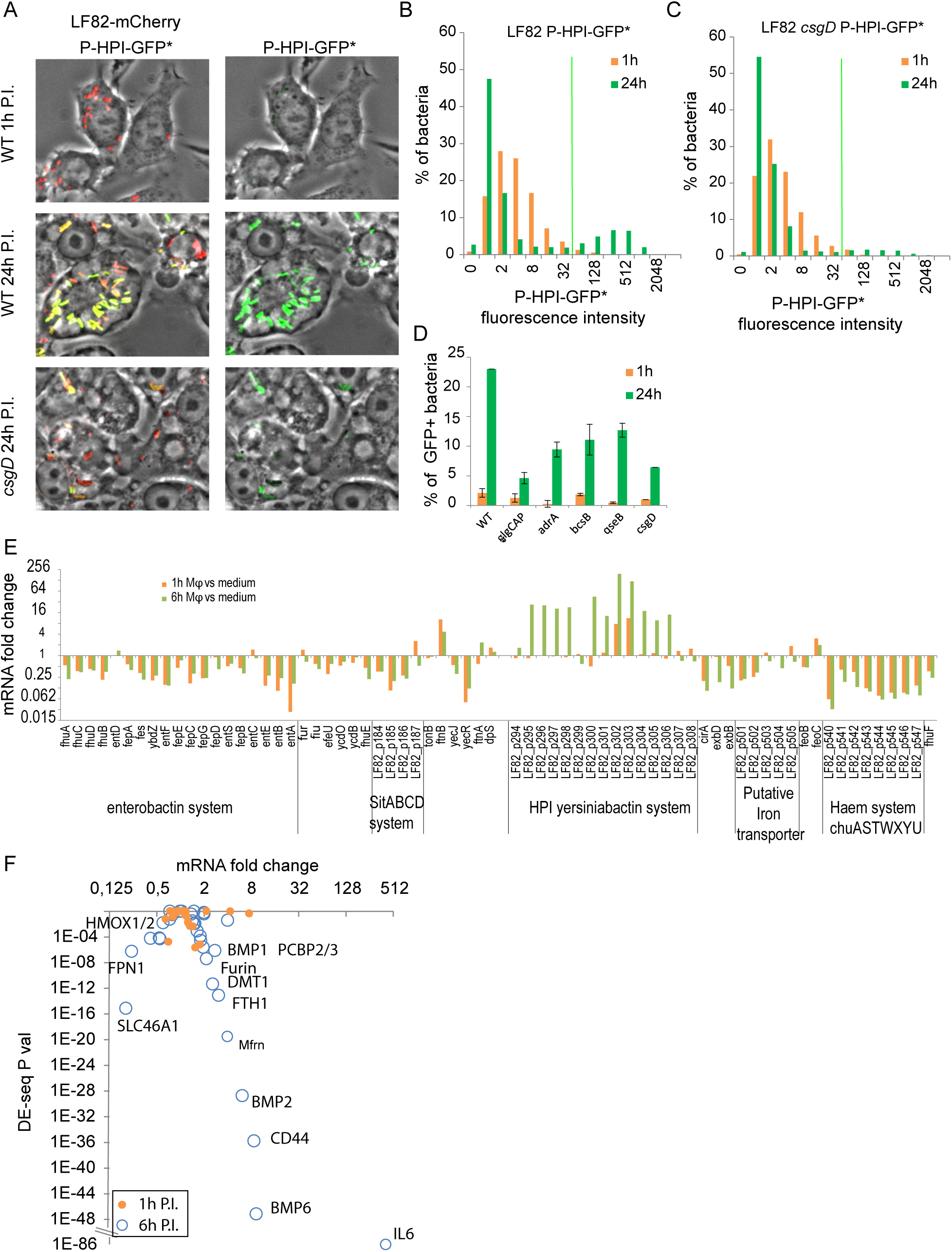
Exopolysaccharide matrix formation and iron homeostasis are interconnected. A) Imaging of the expression of the HPI promoter monitored with an unstable GFP fusion. WT and *csgD* mutant LF82–mCherry strains expressing P-HPI-GFP* from a plasmid were imaged. B) Quantification of the distribution of P-HPI-GFP* fluorescence in LF82. F) Quantification of the distribution of P-HPI-GFP* fluorescence in the LF82 *csgD* mutant. C) Quantification of the % of P-HPI-GFP*-positive cells in the WT strain and the *glgCAP, adrA, bcsB, qseB* and *csgD* LF82 mutants. E) Expression data from RNA-seq experiments at 1 h and 6 h P.I. of THP1 macrophages for the genes annotated for their participation in iron homeostasis in LF82. Genes were organized according to their position on the genome. Clusters of genes are delineated by a vertical line. F) Expression data from RNA-seq experiments at 1 h and 6 h P.I. in THP1 macrophages for the genes annotated for their participation in iron homeostasis in humans. Selected genes with significant expression changes are noted.

### LF82 induces an iron response in the macrophage

We explored human RNA-seq data for eventual signs of iron homeostasis dysregulation (Figure 6F). Interestingly, the expression of several genes involved in iron homeostasis pathway was changed at 6h P.I.. The IL6, ferritin, iron chaperones PCBP 1-3 and the ferrous iron transporter DMT1 were significantly overexpressed, while we noticed a reduction in expression of ferroportin, HCP1, HMOX1 and HMOX2. Surprisingly, we also noted that HAMP and LCN2 genes, encoding hepcidin and lipocalin, respectively, two of the main players of hypoferremia in response to infection by other pathogens, remained unaffected after infection (Table S7). These data support a modification of iron homeostasis in the macrophage in correlation with the formation of LF82 IBCs, such synergy might reflect the expulsion of iron from phagolysosomes and its retention in ferritin within the cytoplasm or mitochondria of macrophages.

## Discussion

### Within phagolysosomes, LF82 presents the characteristics of an IBC

Although there is a strong line of evidence supporting the role of AIEC in promoting gut inflammation and exacerbating CD pathology, the genetic determinants that discriminate AIEC from commensals have remained elusive. In the present work, we demonstrated that LF82 forms IBCs with biofilm-like traits inside phagolysosomes. Immediately upon infection, stringent response induction slowed the LF82 growth rate for a 6 to 10h period. Late during this step, we observed the induction in bacteria of many genes involved in the biofilm pathway (Figure 2). At 10h, LF82 growth resumed concomitantly with the appearance of an extracellular matrix composed of exopolysaccharides and curli fibers around the bacteria (Figure 3). Deletion (Figure 4), interruption by a transposon of genes involved in the biofilm pathway (Figure 5) or chemical inhibition by cis-2-decenoic acid (Figure 4) stopped the formation of the exopolysaccharide matrix and was detrimental for LF82 survival within murine and human macrophages (Figure 4). Altogether, these results suggest that IBCs are an integral part of the adaptation that LF82 must undergo to survive within phagolysosomes. IBCs with biofilm-like traits have also been observed for UPEC during invasion of the urothelial cells lining the urinary bladder (Anderson *et al*, 2003). In this biofilm state, UPEC evade host defense and persist despite antibiotic therapy. Interestingly, AIEC are genetically closely related to UPEC (Nash *et al*, 2010). After invasion, UPEC replicate rapidly (doubling times of 30-35 minutes) for up to 8 hours and form a loose collection of hundreds of bacteria (Scott *et al*, 2015). At this point, the growth rate slows down, and the UPEC develop pods with biofilm-like traits. The kinetics of UPEC’s IBC formation differ from those observed for LF82; the cells remain in a non-replicative state for several hours P.I., and the biofilm traits appear concomitantly with the return to growth of a subpopulation. We also observed an exopolysaccharide matrix inside phagolysosomes containing few (2 to 5) LF82 bacteria (Figure 3). This suggests that bacterial multiplication occurs inside the biofilm for IBC development.

### What are the selective advantages conferred by the IBC?

Biofilms formed on abiotic surfaces or on the epithelium are known to provide resistance to mechanical stress exerted by flow. At first glance, this might not be an important characteristic for IBCs that are confined within the phagolysosomal membrane. However, this characteristic might play a role in the long term if the macrophages die and release the IBCs. This structure might protect LF82 from dispersal by flow or from antibiotics present in the digestive tract and eventually from phagocytosis by another macrophage. Once phagocytosed by a secondary macrophage, the IBC might provide an immediate important bacterial load that is harder for macrophages to eliminate than single bacteria. In Crohn’s disease patients, such a phenomenon might explain the chronic resurgence of AIEC (Alhagamhmad *et al*, 2016). Alternatively, it is possible that the biofilm acts as a physical barrier against the toxic compounds produced by macrophages, perhaps by limiting their diffusion into the core of the IBCs (Mah & O’Toole, 2001). It has been observed that resistance to antimicrobial peptides is mainly based on the interaction between peptides and biofilm exopolymers. These polymers may work by electrostatic repulsion and/or sequestration of antibacterial substances (Otto, 2006). Finally, we can also postulate that IBCs provide a nutrient niche for LF82. The strong stringent response induction and the metabolic switches that we detected in the RNA-seq data suggest that phagolysosomal activity leads to a severe depletion in nutrients for LF82. Cannibalism of dead LF82 might occur in the phagolysosome, and the matrix may facilitate this process (López *et al*, 2009). During coinfection experiments, we observed K12-C600 *E. coli* within LF82 IBCs (Figure 3). We can postulate that in the context of the human gut, such multispecies IBCs are formed and that LF82 can feed on commensal bacteria. The presence of putative T6SSs in the genome of LF82 suggests that this strain is well armed to kill foreign IBC residents.

### HPI is required by IBCs within phagolysosomes

Using molecular genomic methods, we surveyed the genome of LF82 to identify determinants that allow this bacterium to form structured IBCs while commensal *E. coli* do not form such IBCs. LF82 harbors 925 genes that are absent from the genome of K12 MG1655. Among these genes, only a few seem to be specific to AIEC (Nash *et al*, 2010); therefore, it is likely that the phenotypic characteristics of these bacteria emerged from a combination of genes or from pathoadaptive mutations (Miquel *et al*, 2010b). Here, we present evidence suggesting that LF82 requires the combined action of the biofilm pathway encoded on the *E. coli* core genome and a particular iron capture system encoded by a pathogenicity island (HPI) acquired by horizontal transfer to establish the IBC. An analysis of the genetic diversity of *E. coli* isolated from patients with Crohn’s disease revealed that 25 out of 32 strains (78%) encode the irp2 gene and presumably the rest of the HPI cluster in their genome. Among these 32 strains, only 13 presented an AIEC phenotype, and the prevalence of *irp2* in this group was 70% (Céspedes *et al*, 2017). In contrast, this association was only 30% for the control group. In addition, this analysis revealed that the presence of *chuA, fhuD* or *irp*2, three genes involved in iron capture systems, in the genome is a strong marker for *E. coli* associated with Crohn’s disease. *Enterobacteriaceae* (*Yersinia* species or extra intestinal pathogenic *E. coli* (ExPEC)) that harbor HPI exhibit increased virulence. The role of HPI is to synthesize, export and capture Yersiniabactin bound to Fe^3+^. Therefore, Yersiniabactin facilitates iron uptake, which is essential for bacterial growth. In an environment with very limited free iron, such as the human body, where most iron is bound to proteins, bacterial expression of the HPI gene has been shown to be elevated (Chauvaux *et al*, 2007). We observed that 6h P.I., HPI was the only iron capture system with increased expression compared to the growth medium. This suggests that Yersiniabactin is particularly well suited for iron capture within macrophages. It has been observed that Yersiniabactin increases *Enterobacter hormaechei* growth when iron-saturated lactoferrin is the main iron source (Paauw *et al*, 2009). Inside THP1 macrophages, transferrin and lactoferrin genes are expressed at a similar level, but lactoferrin is overexpressed 2.5-fold in the presence of LF82 (Figure 6). Additional experiments will be required to test whether Yersiniabactin and lactoferrin are indeed present in the LF82 phagolysosomes and may compete for iron. An alternative possibility could be that the chemical nature of the phagolysosome limits the iron capture propensity of the enterobactin siderophore and other iron transport systems, increasing the selective pressure for Yersiniabactin. Finally, Yersiniabactin might also contribute to the protection of LF82 from iron (Paauw *et al*, 2009) or copper (Koh *et al*, 2017) toxicity, which may be particularly relevant in phagolysosomal compartments and other restricted spaces in which reactive oxygen species are abundant (White *et al*, 2009). Our observations that a moderate dose of Ga^3+^ simultaneously decreased the global viability of LF82 within macrophages and increased the number of IBCs of large size support the hypothesis that IBCs create a more favorable environment than small vacuoles for LF82.

### Iron capture and biofilm formation

Our results show a direct link between iron capture and biofilm formation. We simultaneously observed that HPI expression is controlled by the biofilm pathway and that the formation of the exopolysaccharide matrix is altered when HPI is deleted. This suggests a synergistic effect between these elements. Links between biofilms and iron homeostasis supporting this synergy have already been documented in a few cases. First, it has been observed that the availability of iron greatly influences the ability of UPEC to form biofilms on the abiotic surface in urine medium (a poor source of iron); interestingly, this phenomenon was also dependent on the yersiniabactin system (Hancock *et al*, 2008). Second, *Pseudomonas aeruginosa* uses exopolysaccharides to sequester and store iron and stimulate biofilm formation (Yu *et al*, 2016; Kang & Kirienko, 2018). Negatively charged exopolysaccharides chelate Fe^3+^ and Fe^2+^ in the vicinity of the bacteria and allow its capture. This scenario could be particularly interesting in the closed environment of the phagolysosomes where iron might be expulsed by the DMT1 Fe^2+^ transporter.

### A complex adaptive strategy to colonize a stressful niche

Altogether, our results suggest that for adapting to the harsh environment of the phagolysosomes (Figure 1A and Figure 2B), AIEC use a complex strategy involving a strong transcriptomic response, a phenotypic switch, mechanisms to capture essential nutrients and a molecular shield to protect the IBC from environmental stress. The nutrient scarcity of the phagolysosomes, because it induces a stringent response and HPI expression, appeared to be an essential determinant of this strategy. Future work will be required to determine whether the phenotypic switch (growing–non-growing), production of the extracellular matrix and iron capture are connected pathways. Links between the stringent response and biofilm formation have already been documented in Salmonella (Azriel *et al*, 2015). Common regulators such as DksA, CsgD and c-di-GMP for these three pathways might explain the kinetics of macrophage infection by LF82. Our observations suggest that other pathobiont bacteria equipped with a biofilm production apparatus and an adequate iron capture system might also survive and form IBCs within macrophages. Does the colonization of this difficult niche represent a selective advantage for AIEC or other bacteria in the context of Crohn’s disease or other diseases involving dysbiosis? Investigating this aspect should provide leads for future antibacterial-based strategies for precision medicine.

## Acknowledgements

This work has benefited from the facilities and expertise of the high throughput sequencing core facilities of I2BC (Centre de Recherche de Gif – http://www.i2bc.paris-saclay.fr/) and Curie Institute (supported by the ANR-10-EQPX-03 and ANR10-INBS-09-08 grants from the Agence Nationale de la Recherche (investissements d’avenir) and by the Canceropôle Ile-de-France). We are very grateful to the members of the CIRB imaging facility. We are grateful to Ugo Szachnowski for his help with RNA-seq data processing and Nicolas Lapaque and Eric Allemand for helpful suggestions. This work has received support from the program «Investissements d’Avenir » launched by the French Government and implemented by ANR with the references ANR-10-LABX-54 MEMOLIFE (OE) and ANR-11-IDEX-0001-02 PSL* Research University (OE), from the ANR (https://anr.fr) with the reference ANR-18-CE35-0007 (OE), ANR-15-CE12-0007 (AM), from the European Research Council “EpincRNA” (starting) and “DARK” (consolidator) grants (AM) and the support of the association François Aupetit (AFA, https://www.afa.asso.fr) (OE). The funders had no role in study design, data collection and analysis, decision to publish, or preparation of the manuscript.

## Legend of the Supplementary Figures

**Supplementary Figure S1: A)** Analysis of the 500 LF82 genes presenting the most significantly changed mRNA fold at 1 h P.I.. The data for the 6 h P.I. time point were presented on Figure 2. Gene annotations were manually curated to define 22 categories: Acid pH response, Biofilm matrix and regulation, Cell cycle, Central metabolism (carbohydrates, nucleotides, amino acids), Chemotaxis, Efflux pumps, Energy and respiratory metabolism, Import and export of nutrients, LF82 genes without homologue in the E. coli K12 genome, LF82 genes encoded by the mega plasmid pLF82, Membrane and envelope components, Metal homeostasis, oxidative stress, Phosphate homeostasis, SOS response, Diverse stress response (including cold shock proteins and phage shock proteins), Stringent response, Transcription factors, uncharacterized genes and putative virulence factors (presenting an homolog in the K12 genome). Numbers on the graph indicate the number of genes in the given category. **B)** Top panel, RNA-seq data from LF82 infecting THP1 macrophages for 6 h P.I. compared to liquid medium culture were analyzed according to regulon information. Regulon annotations were extracted from RegulonDB. For each transcription factor, the RNA-seq data from the regulated genes were collected, and only genes with a significant fold change (DEseq P-value < 10^−10^) were considered. The box plot represents the median fold change of upregulated and downregulated genes from each regulon (bar), the distribution of 75% of the population (box) and outliers (cross). Bottom panel, RNA-seq data from LF82 infecting THP1 macrophages for 6 h P.I. compared to liquid medium culture were analyzed according to regulon information. The numbers of upregulated and downregulated genes from each regulon were plotted. **C)** Histogram showing the fold change in the expression of each LF82 gene belonging to the acidic pH response; biofilm and adhesins; flagella; glycolysis and ATP production annotation groups. For each gene, unfiltered RNA-seq data obtained at 1 h P.I. and 6 h P.I. were plotted.

**Supplementary Figure S2: Exopolysaccharide matrix is observed in three different types of macrophages** A) SBA labeling of the LF82 IBC at 1 h, 6 h and 24 h P.I. in Raw 264.7 macrophages. B) WGA labeling of the LF82 IBC at 1 h, 6 h and 24 h P.I. in HDMM macrophages. C) WGA labeling of the LF82 IBC in THP1 macrophages at 1 h and 24 h P.I. D) WGA labeling of the LF82 IBC in Raw 264.7 macrophages at 1 h and 24 h P.I. E) WGA labeling of the LF82 and LF82*csgD* IBC in Raw 264.7 macrophages at 72 h P.I.

**Supplementary Figure S3: LF82 IBC did not present concanavalin A or Peanut Agglutinin (PNA) staining.** A) Concanavalin A labeling of Raw 264.7 macrophages infected or not by LF82. B) PNA labeling of Raw 264.7 macrophages infected by LF82.

**Supplementary Figure S4: In vitro biofilm formation by LF82 and mutants on abiotic surfaces. A) Measure of the** Specific Biofilm Formation (*SBF*) indices by coloration of the wells of polystyrene microplate with crystal violet. LF82 WT and mutants were incubated for 24h at 37°C without agitation before washing and staining. B) Macrocolonies formation by LF82 WT, LF82 mutants and commensal E. coli at ph7.4, 5.5 and 4.7 at 37°C and 25°C

**Supplementary Figure S5: Tn-seq and RNA-seq results for different LF82 gene clusters.**

**Supplementary Figure S6: Live imaging of LF82 mCherry P-HPI-GFP* expression during infection.** A) Snapshot of the field of view at 30 min and 1450 min P.I., B) Montage on one selected macrophage presenting P-HPI-GFP* expression of a LF82’s IBC. C) Montage on one selected macrophage that did not present P-HPI-GFP* expression of a LF82’s bacteria. D) Quantification of P-HPI-GFP* expression over the infection kinetics. The green curve is the average of 10 IBC where visible P-HPI-GFP* expression was detected. The red-orange curves are examples of macrophage where no expression of P-HPI-GFP* was visible.

## Supplementary Tables

**Table S1: RNA-seq alignment data**

**Table S2: RNA-seq data for the LF82 genome**

**Table S3: GO analysis of LF82 RNA-seq data**

**Table S4: Tn-seq data for the LF82 genome**

**Table S5: Tn-seq data and GO analysis of the important and detrimental LF82 genes after 3 macrophage infections**

**Table S6: RNA-seq data for the human genome**

**Table S7: RNA-seq data for iron homeostasis genes in the human genome**

## STAR+METHODS

### KEY RESOURCES TABLE

#### Antibodies

| Name | Source | Identifier |
| --- | --- | --- |
| Anti-LAMP1 antibody | abcam | AB25630 |
| Anti-LAMP1 antibody | abcam | AB25245 |
| CsgA | Gift from Matthew Chapman | (Zhou *et al*, 2013) |

#### Bacterial Strains

| Name | Genotype | Reference |
| --- | --- | --- |
| AIEC LF82 |  | (Glasser *et al.*, 2001) |
| AIEC LF82 Δbla | *ampC* | Gift from Nicolas Barnich |
| C600 |  | Lab stock |
| AIEC LFVP74 | *csgD::kan* | This work |
| AIEC LFVP80 | *glgC-A-P::kan* | This work |
| AIEC LFVP81 | rcsC-B-D::kan | This work |
| AIEC LFVP86 | rseA-B::kan | This work |
| AIEC LFVP87 | waaW-V-L::kan | This work |
| AIEC LFVP88 | bcsB::kan | This work |
| AIEC LFVP89 | fimE::kan | This work |
| AIEC LFVP90 | adrA::kan | This work |
| AIEC LFVP91 | qseB-C::kan | This work |
| AIEC LFVP92 | pgaA::kan | This work |
| AIEC LFVP93 | gmr::kan | This work |
| AIEC LFVP94 | wza::kan | This work |
| AIEC LFVP95 | dgcE::kan | This work |
| AIEC LFAR01 | ybtS-X-Q-P-A-irp1::kan | This work |

#### Plasmids

| Name | Description | Antibiotic resistance | Reference |
| --- | --- | --- | --- |
| pKOBEGA | Recombineering vector | ampR specR | (Derbise *et al.*, 2003) |
| pPrpsm-mcherry | pGBM2-PrpsM-mCherry | specR | This work |
| pOM1-GFP | pGBM2-Pro3-GFP | specR | (Espéli *et al.*, 2001) |
| p1690-P-HPI-GFP* | ybtA and the promoter of irp1 colend<br>SacI –XbaI in front of unstable GFP of<br>the pSM1690 (Sternberg *et al.*, 1999) | kanR | This work |

#### Chemicals, Peptides, and Recombinant Proteins

| Name | Source | Identifier |
| --- | --- | --- |
| FluoProbes 647H - Phalloidin (653/675nm) | Interchim | FP-BZ9630 |
| FluoProbes 405 Phalloidin | Interchim | FP-1G6270 |
| WGA-alexafluor 647 conjugate | Thermofisher | W32466 |
| WGA-alexafluor 555 conjugate | Invitrogen | W32464 |
| Concanavalin A, Alexa Fluor™ 647 Conjugate | Thermofisher | C21421 |
| Lectin PNA From Arachis hypogaea (peanut), Alexa Fluor™ 594 Conjugate | Thermofisher | L32459 |
| Lectin SBA From Glycine max (soybean), Alexa Fluor™ 647 Conjugate | Thermofisher | L32463 |
| Cis2 decenoic acid | Coger | CDX-D0249-M100 |
| gallium nitrate | SIGMA | 289892 |
| rh-M-CSF | Immunotools | 11343112 |

#### Experimental Models: Cells and Cell lines

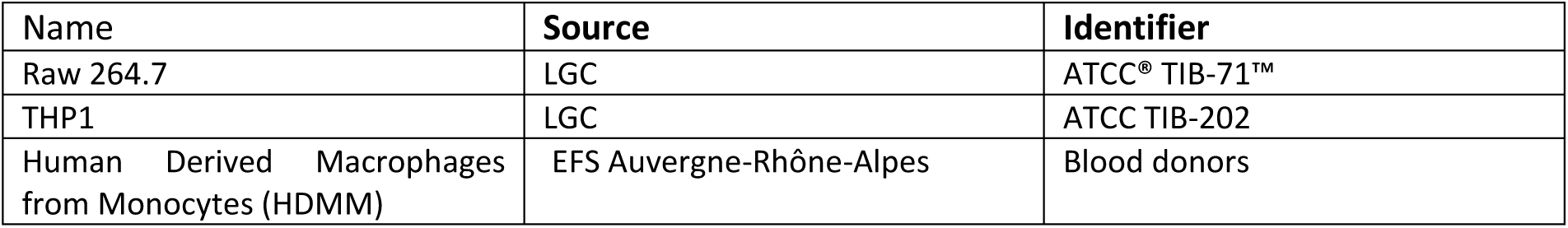

#### Software

| Name | Source | Reference |
| --- | --- | --- |
| Matlab | https://fr.mathworks.com/ |  |
| Tn-seq explorer | https://github.com/sina-cb/Tn-seqExplorer | (Solaimanpour *et al*, 2015) |
| Fiji | https://imagej.net/Fiji |  |
| Cytoscape | https://cytoscape.org/ | (Shannon *et al*, 2003) |

### LEAD CONTACT AND MATERIALS AVAILABILITY

Further information and requests for resources and reagents should be directed to and will be fulfilled by the Lead Contact, Olivier Espeli.

### METHOD DETAILS

#### Infection, viable count and confocal microscopy

Macrophages were infected and imaged as previously described (Demarre *et al.*, 2017). Infections were performed at MOI 30 (measured by CFU), resulting in the observation of 3 LF82 bacteria per macrophage on average at 1 h P.I.. For control experiments with the K12-C600 strain infection were performed at MOI 200. THP1 (ATCC^®^ TIB-202) monocytes (5×10^5^ cells/ml) were differentiated into macrophages for 18 h in phorbol 12-myristate 13-acetate (PMA, 20 ng/ml) before infection. Peripheral blood monocyte-derived macrophages (MDM) were obtained from blood donors as previously described (Vazeille *et al*, 2015; Buisson *et al*, 2019). Briefly, monocytes were purified and differentiated into MDM for 6 days in presence of 0.2μg/ml of recombinant human macrophage colony stimulating factor (rh-M-CSF, Immunotools, Friesoythe, Germany).

Antibiotic challenge and viable bacterial count was performed as described before (Demarre *et al*, 2019). For imaging, cells were cultured on a glass microscope coverslip at the bottom of the microplate wells. Cell were fixed in fresh paraformaldehyde buffer (1X PBS, 3.7% paraformaldehyde) for 40 min and then washed 3 times in 1X PBS. For direct immunostaining, fixed cells were permeabilized in permeabilization buffer (0.05% saponin, 10% Veal fetal Serum, 1X PBS). Lamp1 antibody was added at a 1/200 dilution for 1 h. Coverslips were washed three times in permeabilization buffer and the secondary antibody was added at a 1/1000 dilution for 1 h. When it is required, fluorescent lectin was added at the same time as the secondary antibody. Cover slips were washed 5 times in 1X PBS, dried and mounted in Dako S3023. Imaging was performed on an inverted Zeiss Axio Imager with a spinning disk CSU W1 (Yokogawa). To monitor the presence of curli the macrophage were lysed in lysis buffer (1% Triton, 1X PBS), bacteria were collected by centrifugation for 1 min at 3500g and immediately fixed in paraformaldehyde buffer. Bacteria were layered on polylysine glass coverslips. CsgA antibody (1/1000 dilution) was incubated for 1 h in 1X PBS, washed 3 times in 1X PBS and the secondary antibody coupled to Alexa 645 was added for 1 h. Imaging was performed on an inverted Zeiss Axio Imager with a spinning disk CSU W1 (Yokogawa) equipped with an incubation chamber (Zeiss) and an Orca Flash camera.

#### FRAP

FRAP was performed on Raw 264.7 macrophages infected with LF82 –GFP. Infection was performed in fluorodish (World Precision Instruments). FRAP was performed at 24h P.I.. Images were acquired using a Plan-APO 60x/1.4NA objective on a Ti Nikon microscope enclosed in a thermostatic chamber (Life Imaging Service) equipped with a Evolve EMCCD camera coupled to a Yokogawa CSU-X1 spinning disk. Metamorph Software (Universal Imaging) was used to collect data.

#### Live imaging

Live imaging was performed on Raw 264.7 macrophages infected with LF82 –mCherry pP-HPI-GFP*. Infection was performed in fluorodish (World Precision Instruments). Imaging was performed on an inverted Zeiss Axio Imager with a spinning disk CSU W1 (Yokogawa) equipped with an incubation chamber (Zeiss) and an Orca Flash camera.

#### STED imaging

Infected macrophage were fixed and immunostained as described above. Imaging was performed with a STED expert Line Abberior Instruments GmbH coupled to a SliceScope Scientifica microscope. Images were acquired in two steps, first the confocal configuration for the mCherry bacteria (561 nm) and the Lamp1 staining (485 nm) and second with the STED configuration for the WGA staining (excitation at 640 nm, depletion at 775 nm). Detection was performed with avalanche photodiodes with filter cubes for green (500-550 nm), red (605-625 nm) and far red (650-720nm) fluorescence.

#### RNA-seq

Total RNA were extracted from THP1 macrophages infected by WT LF82 at 1 h and 6h P.I. Control experiments were performed with macrophages alone and bacteria alone cultivated for 1 h and 6 h in DMEM medium at 37°C and 5% CO_2_ without agitation. RNA extraction was performed as described before (Demarre et al, 2019). Experiments were performed from biological duplicate. Human rRNAs were depleted using the Ribo-Zero kit, then strand-specific libraries were constructed using the TruSeq Stranded Total RNA kit (Illumina). Paired-end sequencing (2 × 100 nt) was performed on a HiSeq 25000 sequencer. Reads alignment was performed using Bowtie 2 (Langmead & Salzberg, 2012) and significant fold changes accessed with the DE-seq pipeline (Anders & Huber, 2010). Fastq files were mapped with TopHat (v2.0.6), once on LF82 reference genome, with intron size limited to 5-20nt (options -i 5 & -I 20), and once on human reference genome hg19. Only uniquely mapping reads were kept (option -g 1). Count table for THP1 data were obtained with HTSeq, and count table for LF82 data were obtained using a custom R script, were each gene count is incremented when overlapped by a fragment. Alignment data are available on Table S1. RNA seq data for LF82 genome are available on Table S2 and on Table S6 for the Human genome.

#### Tn-seq

Tn-seq was performed as described (Yamaichi & Dörr, 2017) with the exception that Mariner transposon’s library was generated by electrotransformation of LF82 with the pTSC189Mariner vector instead of conjugation that is not efficient with LF82. About 2 million independent clones, corresponding to 20 electrotransformations, were collected. The library is diluted to an OD of 0.5 and 6 × 150 µl (≈ 4 × 10^7^ bacteria) were used to infect ≈ 6 × 10^6^ Raw 264.7 macrophages cultured in 6 × 2ml wells. After 24 h of infection macrophage were lysis buffer, bacteria were pelleted. Genomic DNA was immediately extracted from 2 wells and pulled together; this sample corresponds to the first round of selection (G1). Bacteria from the other wells were transferred to 100 ml LB flask and grow overnight at 37°C. Bacterial cultures were diluted to an OD of 0.5 and used to infect 4 wells of Raw 264.7 macrophages as previously. After 24 h of infection macrophage were lysis buffer, bacteria were pelleted. Genomic DNA was immediately extracted from 2 wells and pulled together; this sample corresponds to the second round of selection (G2). Bacteria from the last 2 wells were transferred to 100 ml LB flask and processed as described previously for the third round of selection (G3). In parallel the same library was cultured for 3 rounds in LB flask to access the selective pressure imposed by successive in vitro LB cultures. Genomic DNA was extracted at each step and processed for illumina sequencing as described in (Yamaichi & Dörr, 2017). Sequencing was performed at the Imagif sequencing facility on a miSeq Illumina sequencer. Sequencing data were aligned and analyzed with the Tn-seq explorer software (Solaimanpour *et al*, 2015). Tn-seq data are available on Table S4. The whole Tn-seq process, including the initial library preparation, was performed twice. We normalized the number of insertions sequenced for each experiment. At the genome level, we did not observe a statistically significant difference between replicates 1 and 2 (Pearson correlation = 0.79 - 0.88); therefore, we used the average of replicates 1 and 2 as indicative values.

#### In vitro Biofilm assays

Overnight cultures of LF82, LF82 mutants or K12 C600 E. coli were diluted 1/100 and 150µl were transferred to 96 wells polystyrene plates. Plates were incubated for 24h at 37°C without agitation. OD_600nm_ of each well was recorded. Non adhesive bacteria were discarded. Biofilms were washed twice in 1x PBS and fixed with Bouin buffer (Sigma) for 1h at 60°C. Biofilms were washed twice in 1x PBS and stained with crystal violet for 20 min. Biofilms were washed twice in 1x PBS. Biofilm stained with crystal violet were eluted with 95% Ethanol for 30 min and the absorbance measured at OD_570nm_ with a Tecan microplate reader. Biofilm formation is estimated by the SBF factor (Specific Biofilm Formation): SBF = (OD_570nm_ sample - OD_570nm_ control)/ OD_600nm_. Data are average of three replicates +/- standard deviation.

#### Macrocolony formation

### QUANTIFICATION AND STATISTICAL ANALYSIS

Data were consistently reproduced in at least 3 independent experiments (except for RNA-seq and Tn-seq) All statistics were calculated in Excel software using two-tailed Mann-Whitney tests and are noted in figure legends. P values are expressed as follows: * p < 0.05, ** p < 0.01, *** p < 0.001, **** p < 0.0001.

#### DATA AND CODE AVAILABILITY

RNA-seq and Tn-seq data are available at GEO

## Bibliography

Alhagamhmad MH, Day AS, Lemberg DA & Leach ST (2016) An overview of the bacterial contribution to Crohn disease pathogenesis. J. Med. Microbiol. 65: 1049–1059

Anderson GG, Palermo JJ, Schilling JD, Roth R, Heuser J & Hultgren SJ (2003) Intracellular bacterial biofilm-like pods in urinary tract infections. Science 301: 105–107

Anders S, Huber W (2010). “Differential expression analysis for sequence count data.” Genome Biology, 11, R106.

Anisimov R, Brem D, Heesemann J & Rakin A (2005) Molecular mechanism of YbtA-mediated transcriptional regulation of divergent overlapping promoters ybtA and irp6 of Yersinia enterocolitica. FEMS Microbiol. Lett. 250: 27–32

Atri C, Guerfali FZ, Laouini D. (2018) Role of Human Macrophage Polarization in Inflammation during Infectious Diseases. J Mol Sci. 19(6):1801

Azriel S, Goren A, Rahav G & Gal-Mor O (2015) The Stringent Response Regulator DksA Is Required for Salmonella enterica Serovar Typhimurium Growth in Minimal Medium, Motility, Biofilm Formation, and Intestinal Colonization. Infect. Immun. 84: 375–384

Buisson A, Douadi C, Ouchchane L, Goutte M, Hugot JP, Dubois A, Minet-Quinard R, Bouvier D, Bommelaer G, Vazeille E, Barnich N (2019) Macrophages Inability to Mediate Adherent-Invasive E. coli Replication is Linked to Autophagy in Crohn’s Disease Patients. Cells. 2019 Nov 5;8(11):1394

Bringer M-A, Barnich N, Glasser A-L, Bardot O & Darfeuille-Michaud A (2005) HtrA stress protein is involved in intramacrophagic replication of adherent and invasive Escherichia coli strain LF82 isolated from a patient with Crohn’s disease. Infect. Immun. 73: 712–721

Bringer M-A, Glasser A-L, Tung C-H, Méresse S & Darfeuille-Michaud A (2006) The Crohn’s disease-associated adherent-invasive Escherichia coli strain LF82 replicates in mature phagolysosomes within J774 macrophages. Cell. Microbiol. 8: 471–484

Bringer M-A, Rolhion N, Glasser A-L & Darfeuille-Michaud A (2007) The oxidoreductase DsbA plays a key role in the ability of the Crohn’s disease-associated adherent-invasive Escherichia coli strain LF82 to resist macrophage killing. J. Bacteriol. 189: 4860–4871

Céspedes S, Saitz W, Del Canto F, De la Fuente M, Quera R, Hermoso M, Muñoz R, Ginard D, Khorrami S, Girón J, Assar R, Rosselló-Mora R & Vidal RM (2017) Genetic Diversity and Virulence Determinants of Escherichia coli Strains Isolated from Patients with Crohn’s Disease in Spain and Chile. Front. Microbiol. 8: 639

Chauvaux S, Rosso M-L, Frangeul L, Lacroix C, Labarre L, Schiavo A, Marceau M, Dillies M-A, Foulon J, Coppée J-Y, Médigue C, Simonet M & Carniel E (2007) Transcriptome analysis of Yersinia pestis in human plasma: an approach for discovering bacterial genes involved in septicaemic plague. Microbiol. Read. Engl. 153: 3112–3124

Choi S-R, Britigan BE & Narayanasamy P (2019) Iron/Heme Metabolism-Targeted Gallium(III) Nanoparticles Are Active against Extracellular and Intracellular Pseudomonas aeruginosa and Acinetobacter baumannii. Antimicrob. Agents Chemother. 63:

Demarre G, Prudent V & Espéli O (2017) Imaging the Cell Cycle of Pathogen E. coli During Growth in Macrophage. Methods Mol. Biol. Clifton NJ 1624: 227–236

Demarre G, Prudent V, Schenk H, Rousseau E, Bringer M-A, Barnich N, Tran Van Nhieu G, Rimsky S, De Monte S & Espéli O (2019) The Crohn’s disease-associated Escherichia coli strain LF82 relies on SOS and stringent responses to survive, multiply and tolerate antibiotics within macrophages. PLoS Pathog. 15: e1008123

Derbise A, Lesic B, Dacheux D, Ghigo JM & Carniel E (2003) A rapid and simple method for inactivating chromosomal genes in Yersinia. FEMS Immunol. Med. Microbiol. 38: 113–116

Espéli O, Moulin L & Boccard F (2001) Transcription attenuation associated with bacterial repetitive extragenic BIME elements. J. Mol. Biol. 314: 375–386

Flemming H-C & Wingender J (2010) The biofilm matrix. Nat. Rev. Microbiol. 8: 623–633

Gama-Castro S, Salgado H, Santos-Zavaleta A, Ledezma-Tejeida D, Muñiz-Rascado L, García-Sotelo JS, Alquicira-Hernández K, Martínez-Flores I, Pannier L, Castro-Mondragón JA, Medina-Rivera A, Solano-Lira H, Bonavides-Martínez C, Pérez-Rueda E, Alquicira-Hernández S, Porrón-Sotelo L, López-Fuentes A, Hernández-Koutoucheva A, Del Moral-Chávez V, Rinaldi F, et al (2016) RegulonDB version 9.0: high-level integration of gene regulation, coexpression, motif clustering and beyond. Nucleic Acids Res. 44: D133–143

Gao H, Zhou D, Li Y, Guo Z, Han Y, Song Y, Zhai J, Du Z, Wang X, Lu J & Yang R (2008) The iron-responsive Fur regulon in Yersinia pestis. J. Bacteriol. 190: 3063–3075

Glasser AL, Boudeau J, Barnich N, Perruchot MH, Colombel JF & Darfeuille-Michaud A (2001) Adherent invasive Escherichia coli strains from patients with Crohn’s disease survive and replicate within macrophages without inducing host cell death. Infect. Immun. 69: 5529–5537

Hancock V, Ferrières L & Klemm P (2008) The ferric yersiniabactin uptake receptor FyuA is required for efficient biofilm formation by urinary tract infectious Escherichia coli in human urine. Microbiol. Read. Engl. 154: 167–175

Joo H-S & Otto M (2012) Molecular basis of in vivo biofilm formation by bacterial pathogens. Chem. Biol. 19: 1503–1513

Kang D & Kirienko NV (2018) Interdependence between iron acquisition and biofilm formation in Pseudomonas aeruginosa. J. Microbiol. Seoul Korea 56: 449–457

Koh E-I, Robinson AE, Bandara N, Rogers BE & Henderson JP (2017) Copper import in Escherichia coli by the yersiniabactin metallophore system. Nat. Chem. Biol. 13: 1016–1021

Lapaquette P, Bringer M-A & Darfeuille-Michaud A (2012) Defects in autophagy favour adherent-invasive Escherichia coli persistence within macrophages leading to increased pro-inflammatory response. Cell. Microbiol. 14: 791–807

Langmead B, Salzberg S. (2012) Fast gapped-read alignment with Bowtie 2. Nature Methods., 9:357–359.

López D, Vlamakis H, Losick R & Kolter R (2009) Cannibalism enhances biofilm development in Bacillus subtilis. Mol. Microbiol. 74: 609–618

Mah TF & O’Toole GA (2001) Mechanisms of biofilm resistance to antimicrobial agents. Trends Microbiol. 9: 34–39

Miquel S, Claret L, Bonnet R, Dorboz I, Barnich N & Darfeuille-Michaud A (2010a) Role of decreased levels of Fis histone-like protein in Crohn’s disease-associated adherent invasive Escherichia coli LF82 bacteria interacting with intestinal epithelial cells. J. Bacteriol. 192: 1832–1843

Miquel S, Peyretaillade E, Claret L, de Vallée A, Dossat C, Vacherie B, Zineb EH, Segurens B, Barbe V, Sauvanet P, Neut C, Colombel J-F, Medigue C, Mojica FJM, Peyret P, Bonnet R & Darfeuille-Michaud A (2010b) Complete genome sequence of Crohn’s disease-associated adherent-invasive E. coli strain LF82. PloS One 5:

Nash JH, Villegas A, Kropinski AM, Aguilar-Valenzuela R, Konczy P, Mascarenhas M, Ziebell K, Torres AG, Karmali MA & Coombes BK (2010) Genome sequence of adherent-invasive Escherichia coli and comparative genomic analysis with other E. coli pathotypes. BMC Genomics 11: 667

Otto M (2006) Bacterial evasion of antimicrobial peptides by biofilm formation. Curr. Top. Microbiol. Immunol. 306: 251–258

Paauw A, Leverstein-van Hall MA, van Kessel KPM, Verhoef J & Fluit AC (2009) Yersiniabactin reduces the respiratory oxidative stress response of innate immune cells. PloS One 4: e8240

Parker A, Cureoglu S, De Lay N, Majdalani N & Gottesman S (2017) Alternative pathways for Escherichia coli biofilm formation revealed by sRNA overproduction. Mol. Microbiol. 105: 309–325

Scott VCS, Haake DA, Churchill BM, Justice SS & Kim J-H (2015) Intracellular Bacterial Communities: A Potential Etiology for Chronic Lower Urinary Tract Symptoms. Urology 86: 425–431

Shannon P, Markiel A, Ozier O, Baliga NS, Wang JT, Ramage D, Amin N, Schwikowski B & Ideker T (2003) Cytoscape: a software environment for integrated models of biomolecular interaction networks. Genome Res. 13: 2498–2504

Solaimanpour S, Sarmiento F & Mrázek J (2015) Tn-Seq Explorer: A Tool for Analysis of High-Throughput Sequencing Data of Transposon Mutant Libraries. PLOS ONE 10: e0126070

Stapels DAC, Hill PWS, Westermann AJ, Fisher RA, Thurston TL, Saliba AE, Blommestein I, Vogel J, Helaine S (2018) Salmonella persisters undermine host immune defenses during antibiotic treatment. Science. 362(6419):1156–1160.

Sternberg C, Christensen BB, Johansen T, Toftgaard Nielsen A, Andersen JB, Givskov M & Molin S (1999) Distribution of Bacterial Growth Activity in Flow-Chamber Biofilms. Appl. Environ. Microbiol. 65: 4108–4117

Tawfik A, Flanagan PK & Campbell BJ (2014) Escherichia coli-host macrophage interactions in the pathogenesis of inflammatory bowel disease. World J. Gastroenterol. 20: 8751–8763

Vazeille E, Buisson A, Bringer MA, Goutte M, Ouchchane L, Hugot JP, de Vallée A, Barnich N, Bommelaer G, Darfeuille-Michaud A. Monocyte-derived macrophages from Crohn’s disease patients are impaired in the ability to control intracellular adherent-invasive Escherichia coli and exhibit disordered cytokine secretion profile. J Crohns Colitis. 9(5):410–20.

White C, Lee J, Kambe T, Fritsche K & Petris MJ (2009) A role for the ATP7A copper-transporting ATPase in macrophage bactericidal activity. J. Biol. Chem. 284: 33949–33956

Yamaichi Y & Dörr T (2017) Transposon Insertion Site Sequencing for Synthetic Lethal Screening. Methods Mol. Biol. Clifton NJ 1624: 39–49

Yu S, Wei Q, Zhao T, Guo Y & Ma LZ (2016) A Survival Strategy for Pseudomonas aeruginosa That Uses Exopolysaccharides To Sequester and Store Iron To Stimulate Psl-Dependent Biofilm Formation. Appl. Environ. Microbiol. 82: 6403–6413

Zhou Y, Smith DR, Hufnagel DA & Chapman MR (2013) Experimental Manipulation of the Microbial Functional Amyloid Called Curli. In Bacterial Cell Surfaces: Methods and Protocols, Delcour AH (ed) pp 53–75. Totowa, NJ: Humana Press

