## Supplementary figures and images for "The Crohn’s disease-related AIEC strain LF82 assembles a biofilm-like matrix to protect intracellular microcolonies from phagolysosomal attack"

### Supplementary Figure S1

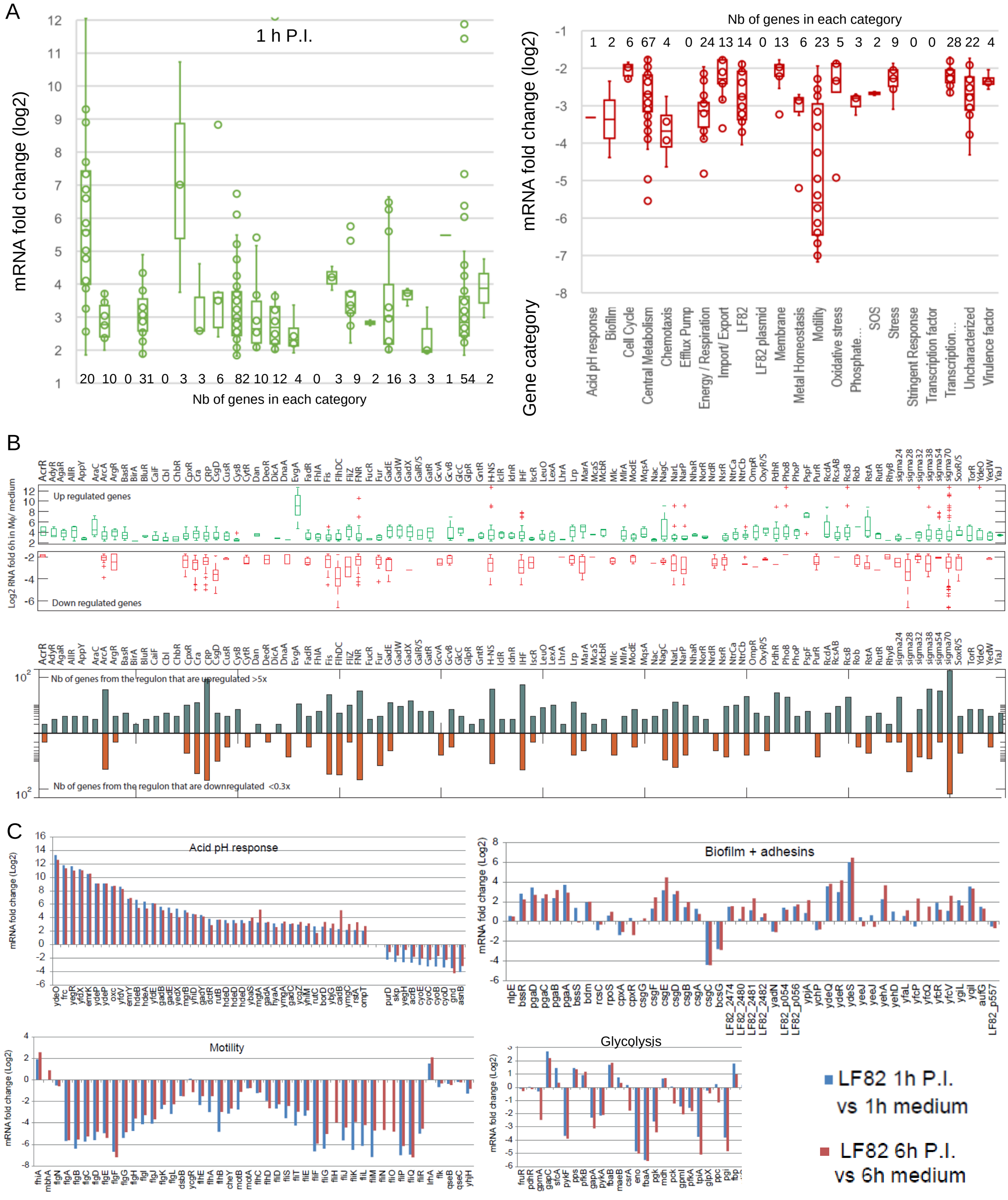

### Supplementary Figure S2

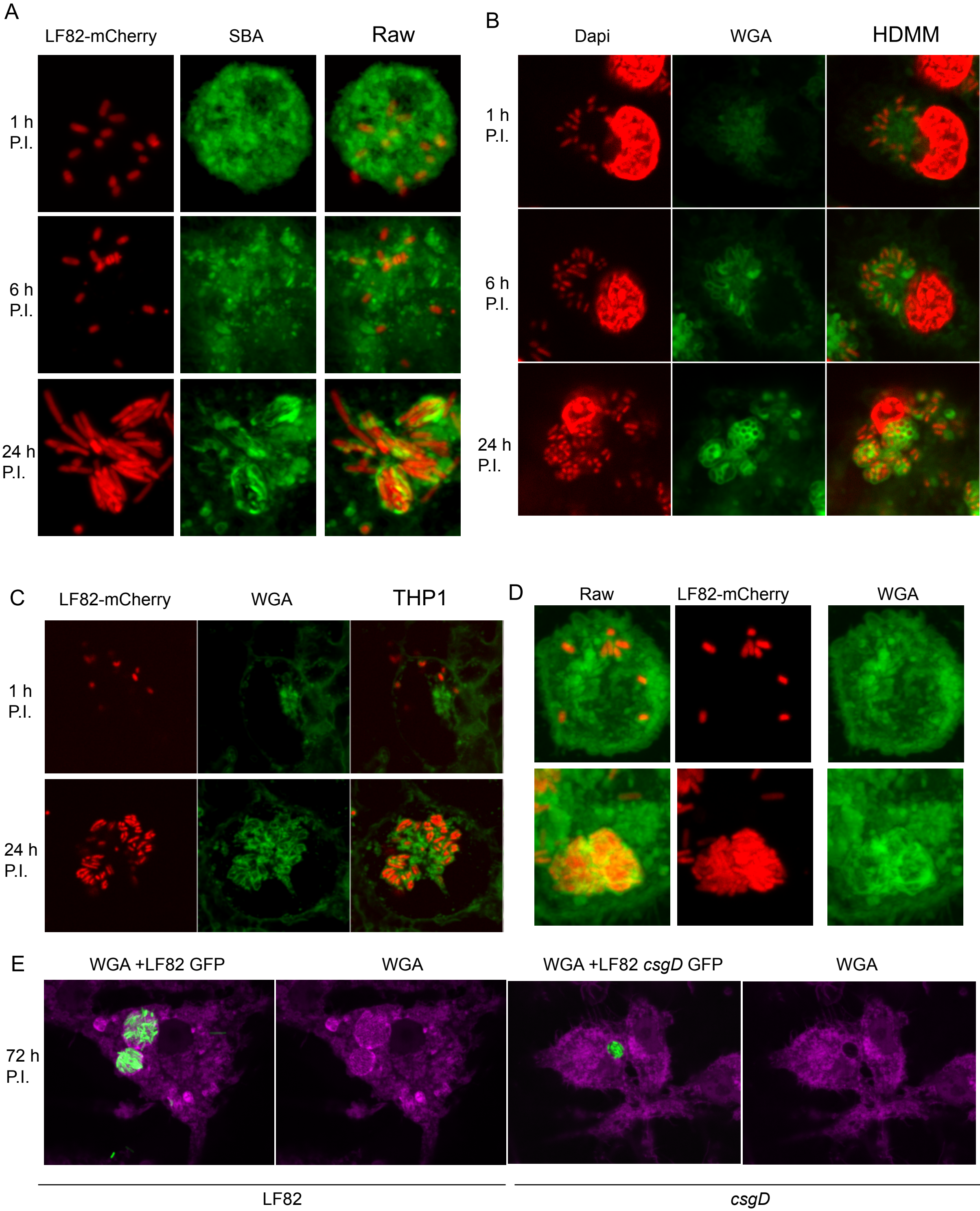

### Supplementary Figure S3

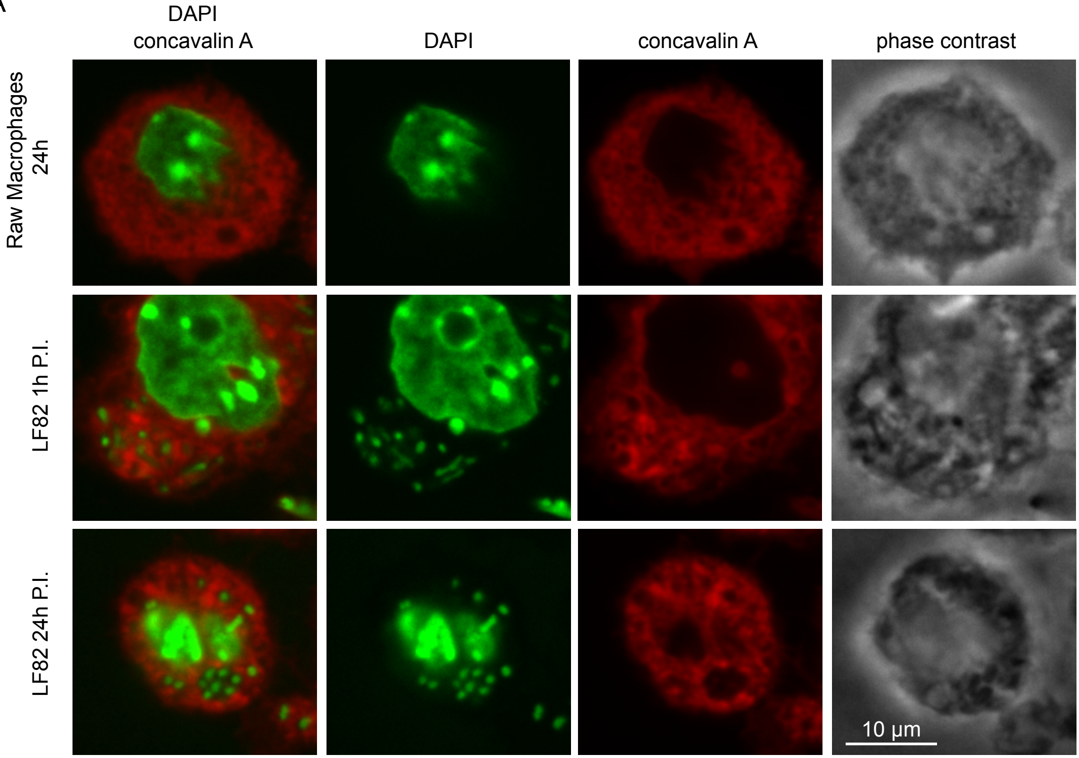

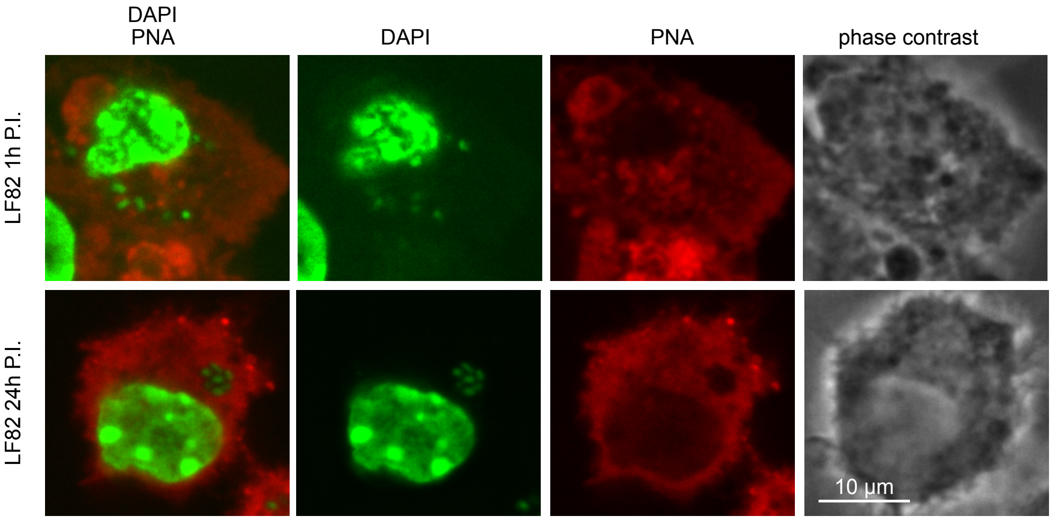

### Supplementary Figure S4

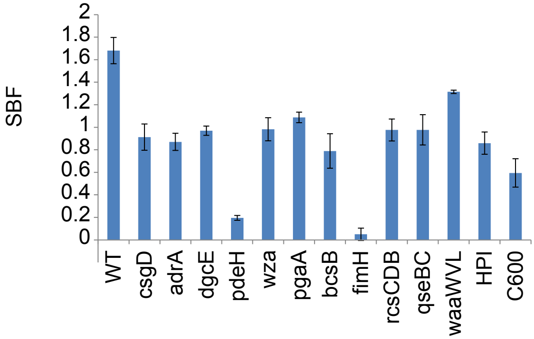

### Supplementary Figure S5

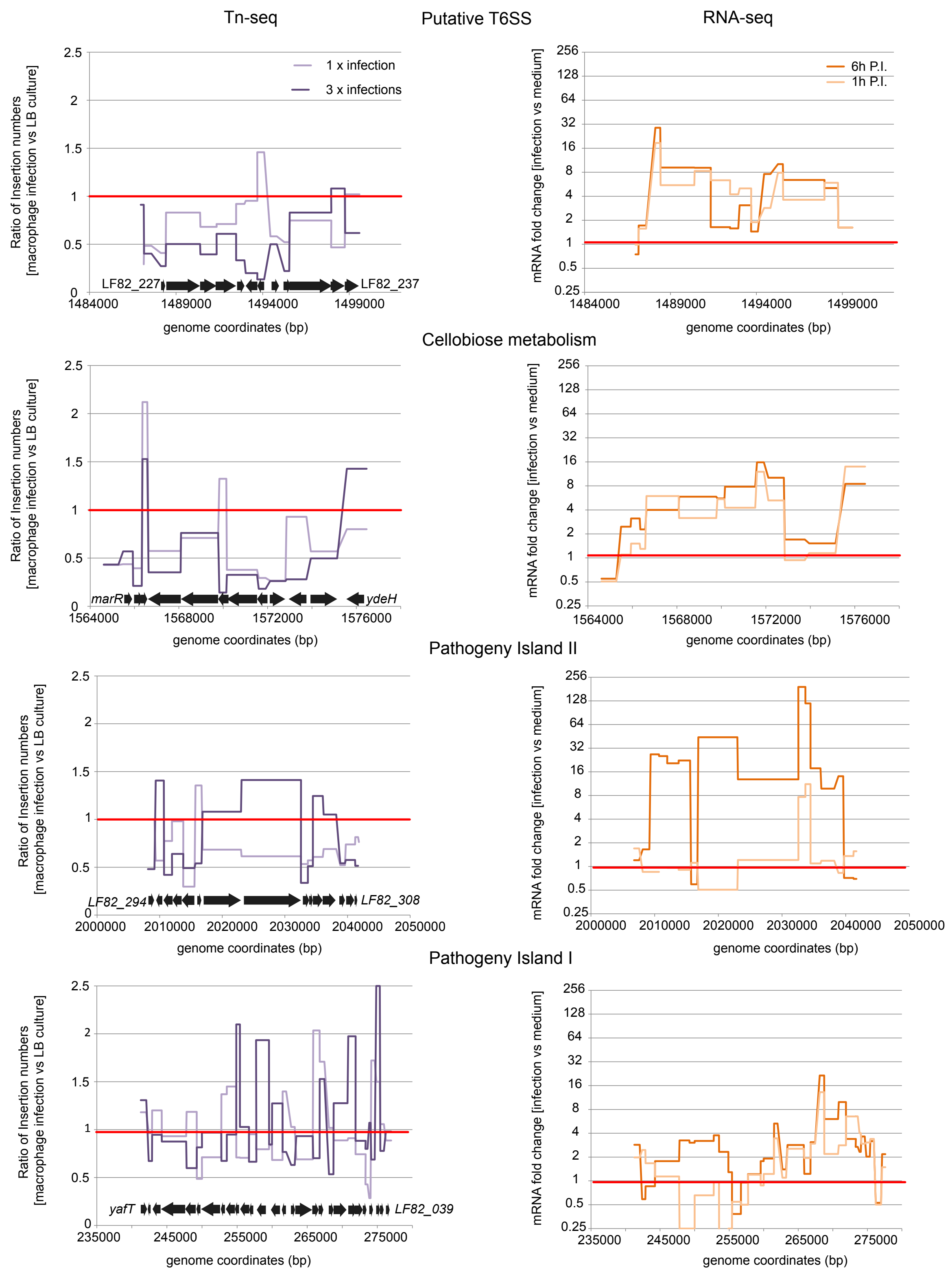

### Supplementary Figure S6

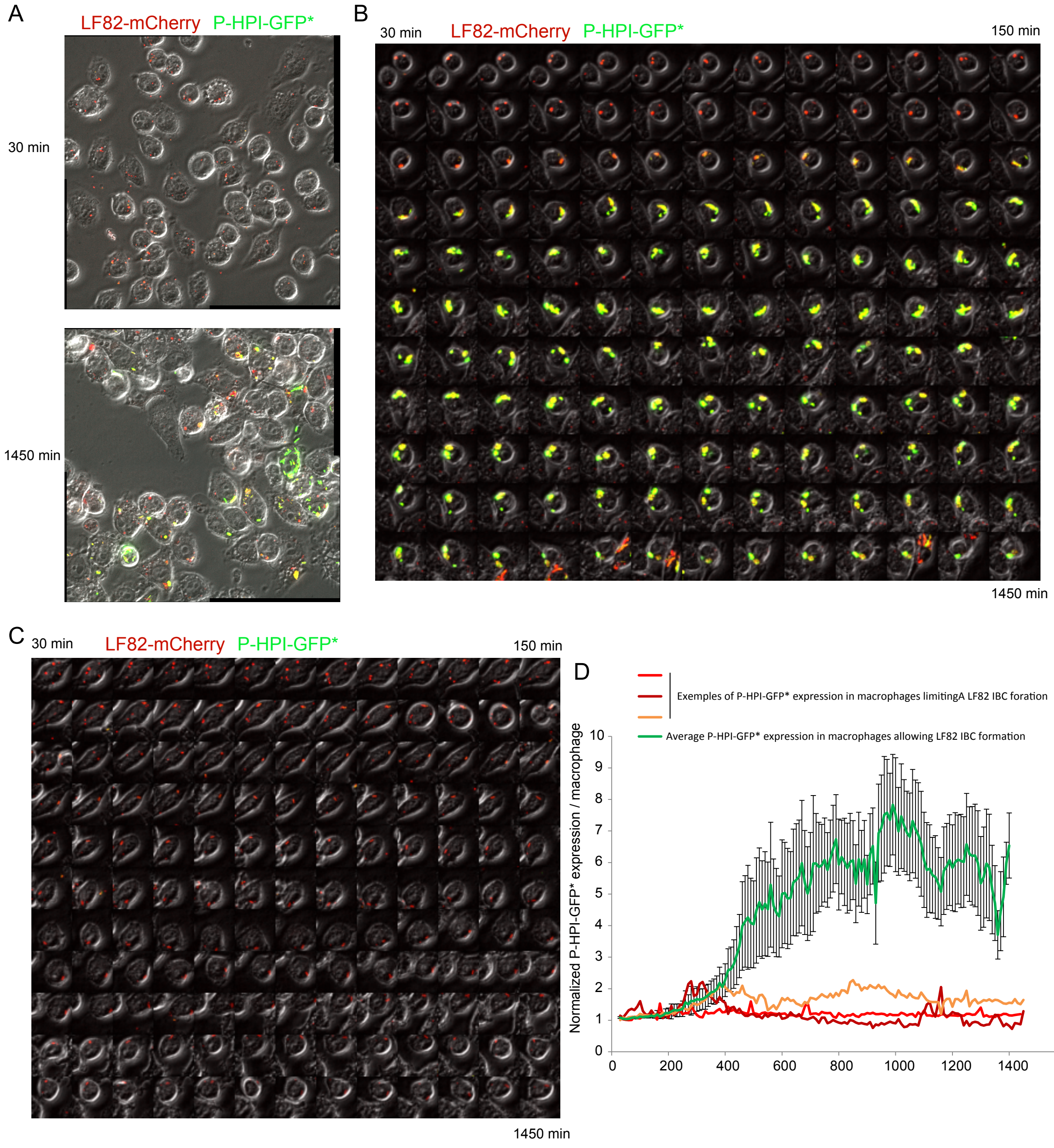
